# Sensory neurons regulate stimulus-dependent humoral immunity

**DOI:** 10.1101/2024.01.04.574231

**Authors:** Diane Aguilar, Fengli Zhu, Antoine Millet, Nicolas Millet, Patrizia Germano, Joseph Pisegna, Omid Akbari, Taylor A. Doherty, Marc Swidergall, Nicholas Jendzjowsky

## Abstract

Sensory neurons sense pathogenic infiltration, informing immune coordination of host defense. However, sensory neuron-immune interactions have been predominantly shown to drive innate immune responses. Humoral memory, whether protective or destructive, is acquired early in life - as demonstrated by both early exposure to streptococci and allergic disease onset. Our study further defines the role of sensory neuron influence on humoral immunity in the lung. Using a murine model of *Streptococcus pneumonia* pre-exposure and infection and a model of allergic asthma, we show that sensory neurons are required for B-cell and plasma cell recruitment and antibody production. In response to *S. pneumoniae*, sensory neuron depletion resulted in a larger bacterial burden, reduced B-cell populations, IgG release, and neutrophil stimulation. Conversely, sensory neuron depletion reduced B-cell populations, IgE, and asthmatic characteristics during allergen-induced airway inflammation. The sensory neuron neuropeptide released within each model differed. With bacterial infection, vasoactive intestinal polypeptide (VIP) was preferentially released, whereas substance P was released in response to asthma. Administration of VIP into sensory neuron-depleted mice suppressed bacterial burden and increased IgG levels, while VIP1R deficiency increased susceptibility to bacterial infection. Sensory neuron-depleted mice treated with substance P increased IgE and asthma, while substance P genetic ablation resulted in blunted IgE, similar to sensory neuron-depleted asthmatic mice. These data demonstrate that the immunogen differentially stimulates sensory neurons to release specific neuropeptides that specifically target B-cells. Targeting sensory neurons may provide an alternate treatment pathway for diseases involved with insufficient and/or aggravated humoral immunity.

**Graphical Abstract:** 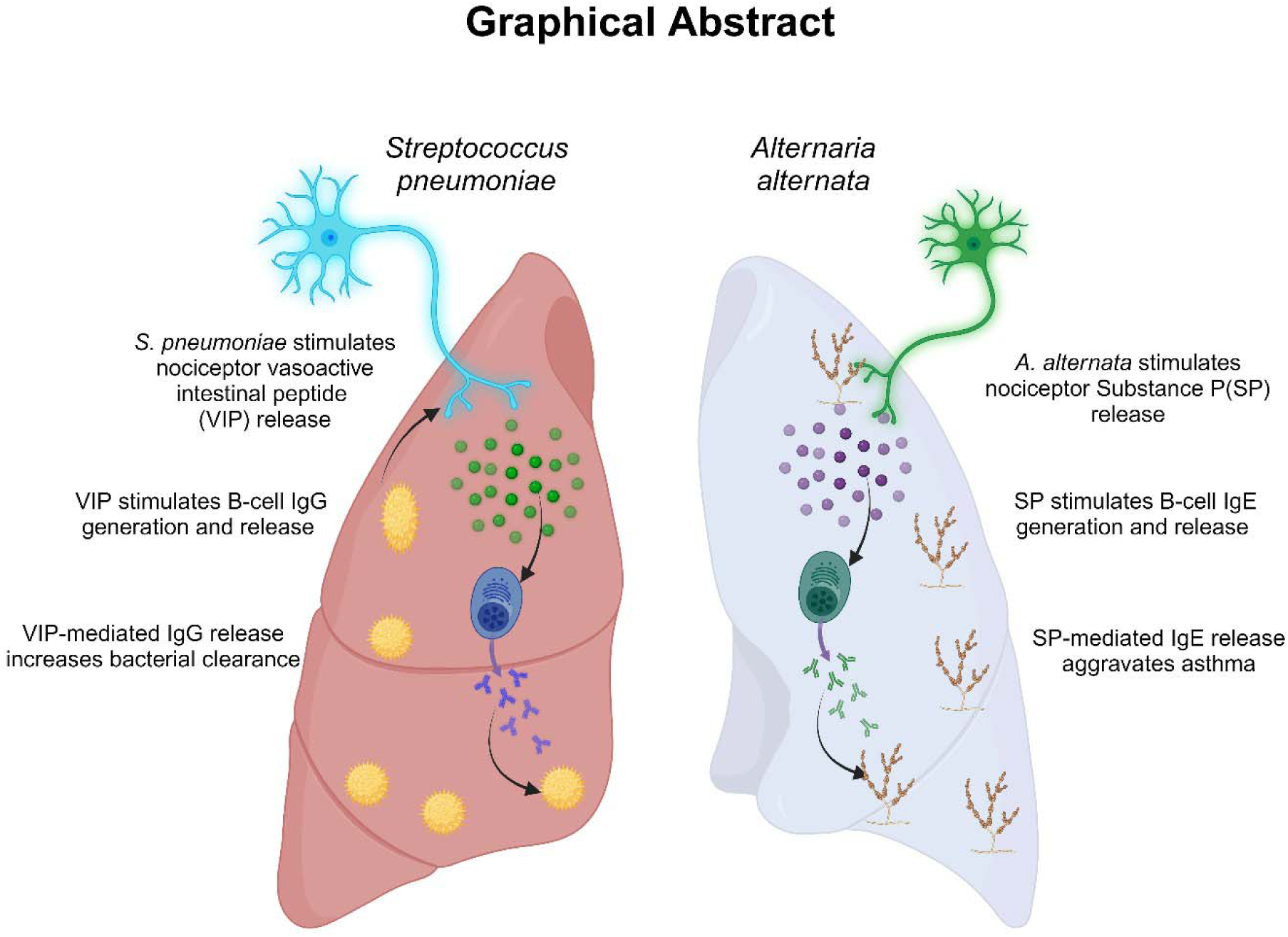

## Introduction

With each breath, the lung risks exposure to atmospheric infectious agents and allergens. Sensory neurons work alongside immune cells and influence host defense against infection and injury^1–7^, but appear to exacerbate allergy^8–12^. Stimulation of sensory neurons by cytokines^4,5,8,12,13^, proteases^11,14–17^, PAMPs^1,1–3^ and immunoglobulins^9,18–20^ lead to neuronal depolarization. The release of neuropeptides by sensory neurons in response to infection or allergy^21–23^, is a necessary cofactor in immune coordination^21,24^. The unique interplay between neurons and immune cells is pathology-dependent. A growing body of work demonstrates immune-stimulus-dependent neuropeptide release and, therefore, differential coordination of varying cell types^1–3,7,25–28^. Such differences speak to the dichotomous neuro-immune response, which can elicit a pro- or anti-inflammatory state^21,29^.

Most current evidence involving sensory neuron and immune cell crosstalk involves cellular-mediated immunity. For example, sensory neurons inhibit bacterial clearance and fungal infection by suppressing Th1 and Th17-mediated cytokine release and, therefore, suppress neutrophil-mediated clearance of infection in response to acute infection in naïve mice^2,7^. Given that many infectious diseases are recurring, prolonged, or stem from commensal microbes becoming invasive^30,31^, further inquiry into the neural influence on humoral immunity in response to infection is warranted.

Conversely, during allergy and asthma, vasoactive intestinal polypeptide (VIP)^8^ and substance P^11^ release, activate ILC2s^8^ and dendritic cells^11^ to increase eosinophilic infiltration^8^ and mast cell degranulation^11^. However, the catalytic event that drives allergic symptoms^32^ is a dysregulated increase of allergen-specific IgE and its downstream effects. Given the very high prevalence of allergic disorders, there is a great need to further understand the role of sensory neurons in regulating humoral immunity, which may also lead to future therapeutic targets.

B-cells are responsible for immune memory by class switch recombination, a process of VDJ recombination where heavy chains are cleaved from IgM to produce IgG, IgA, IgE, or IgD depending on the immune effector response required^33^. Early investigations have shown that B-cells respond to exogenous neuropeptides *in vitro* and can steer the predominant release of specific immunoglobulin^34–38^. For example, B-cells express neurotransmitter receptors for, substance P^34^, vasoactive intestinal peptide^39,40^, noradrenaline, and neuropeptide Y^41,42^. *In vitro*, some of these neurotransmitters appear to increase immunoglobulin release^34–38^. Sensory neurons also mediate B-cell development in the mucosa^39,43^ and promote B-cell class switching in the spleen^44^ and lung^10^. It was shown that sensory neuron ablation reduced IgE release subsequent to a reduction of B220+ cells in the lung, in response to house dust mite asthma and calcipitrol skin allergy^10^. Substance P was demonstrated to be responsible for IgE class switch recombination *in vitro,* and recent *in vivo* results support substance P-mediated class switch^10^. Our study investigates whether sensory neuron neuropeptide release can be a global effector for infection and allergy and whether the interplay between sensory neurons and B-cells is antigen-dependent.

We analyzed how sensory neurons traffic and stimulate B-cells in response to *Streptococcus pneumoniae* pulmonary infection and *Alternaria alternata*-induced allergic airway inflammation. We show that the release of sensory neuropeptides differs between our model of *S. pneumoniae* pre-exposure and infection and in an *A. alternata-*induced asthma model. Sensory neuron depletion suppressed select sensory neuropeptides in the lungs of each model and, as a result, reduced pulmonary B-cell and plasma cell immunoglobulin release. Neuronal depletion also exacerbated infection but improved asthma, proving the necessity of sensory neurons to stimulate humoral immunity. Experiments using neuropeptide knockout mice for respective neuropeptides support neuron depletion data. Supplementation of neuropeptides back into sensory neuron-depleted mice in each model either improved bacterial clearance or worsened asthma features due to a restoration of immunoglobulin production and release- thus supporting the sufficiency of sensory neurons to regulate humoral immunity. In summary, sensory neurons modulate B-cell production of immunoglobulins by variations in neurotransmitter release. The release of neuropeptides is altered by the immune stimulus, which appears to provide a way to steer the humoral response.

## Results

### TRPV1+ neurons mediate survival and bacterial clearance in pneumonia

Transient receptor potential vanilloid 1+ (TRPV1) neurons have been demonstrated to influence T-cell-mediated coordination of immunity by the release of specific neuropeptides^2,3,8^. The TRPV1 ion channel responds to capsaicin, protons, prostaglandins, lipids and heat stimuli^45^. TRPV1+ neurons have been found to coordinate cellular immunity and influence T-cell stimulation of neutrophils in acute *Staphylococcus aureus* infection^2^ and eosinophils and mast cells in allergic asthma^8^. Thus, we tested how TRPV1+ sensory neurons play a role in humoral immunity.

Using an established model of sensory neuron chemical ablation with resiniferatoxin (RTX)^2,46,47^, we suppressed TRPV1-containing neurons similarly in the vagal (nodose/jugular) ganglion by >85% compared to TRPV1-DTR mice injected with the diphtheria toxin directly into the vagal ganglia as assessed by qPCR and TRPV1 immunohistochemistry (**Supplementary** Figure 1). RTX, administered by subcutaneous injection, induces global ablation of TRPV1+ neurons, thus blocking sensory neuron vesicle emission of neuropeptides and rendering them ineffective/atrophied^2,46,47^; we confirm the suppression of neuropeptide transcripts in the vagal ganglia with RTX by qPCR (**Supplementary** Figure 1). Although there are indications that dorsal root ganglia (distal site, primarily innervate skin and visceral organs^48^) provide some innervation of the lungs^49^, recent tracer studies have demonstrated sparse innervation with an overwhelming majority of lung sensory neuron innervation provided by the vagal ganglia^50^.

To investigate humoral immunity in response to *S. pneumoniae* lung infection, we aimed to replicate the colonization and infection patterns observed in humans, considering that mice are not naturally colonized with *S. pneumoniae*^51,52^. We established a model of *S. pneumoniae* infection, which involved exposure of mice with a low inoculum on Day 0, and a subsequent exposure to a high infectious dose of bacteria on Day 9 (**Figure 1a**). Following the low inoculating dose, both sensory neuron intact and RTX mice showed a similar amount of bacteria remaining 16hrs after the low inoculum (**Figure 1b**). Both groups were able to clear the bacteria prior to the infectious dose (**Figure 1b**). Compared to naïve mice, our *S. pneumoniae* model showed an initially increased bacterial burden, which was not different from RTX mice, 16hrs after infection (**Figure 1b**). Forty-eight hours after the last infectious dose of *S. pneumoniae*, sensory neuron intact mice almost completely cleared their bacterial burden, while RTX treated mice exhibited an elevated bacterial burden in comparison to sensory intact mice and naïve mice and this persisted 6 days after the final infection(**Figure 1b**). We further assessed the role of sensory neurons in cross-protection against a lethal pneumococcal strain. Sensory neuron ablation resulted in a higher bacterial burden and reduced survival when pre-exposed to serotype 19F and infected with serotype 3 (**Figure 1c, d**). Additionally, in response to a single high dose of *S. pneumoniae,* in both sensory neuron intact and RTX ablated mice, bacterial burden and IgG remained unchanged, and when exposed to a single infectious dose of serotype 3, survival was unchanged (**Supplementary** Figure 2); similar to previous investigations^2^. Bacterial clearance between RTX-treated and targeted vagal TRPV1 ablation with direct diphtheria toxin (DTX) injection into the vagal ganglia of TRPV1-DT receptor (DTR) mice^53^ was similar (**Supplementary** Figure 3). Consequently, our findings suggest that vagal-specific sensory neurons are required to control *S. pneumoniae* during the course of lung infection in our “recall” model.

**Figure 1.**
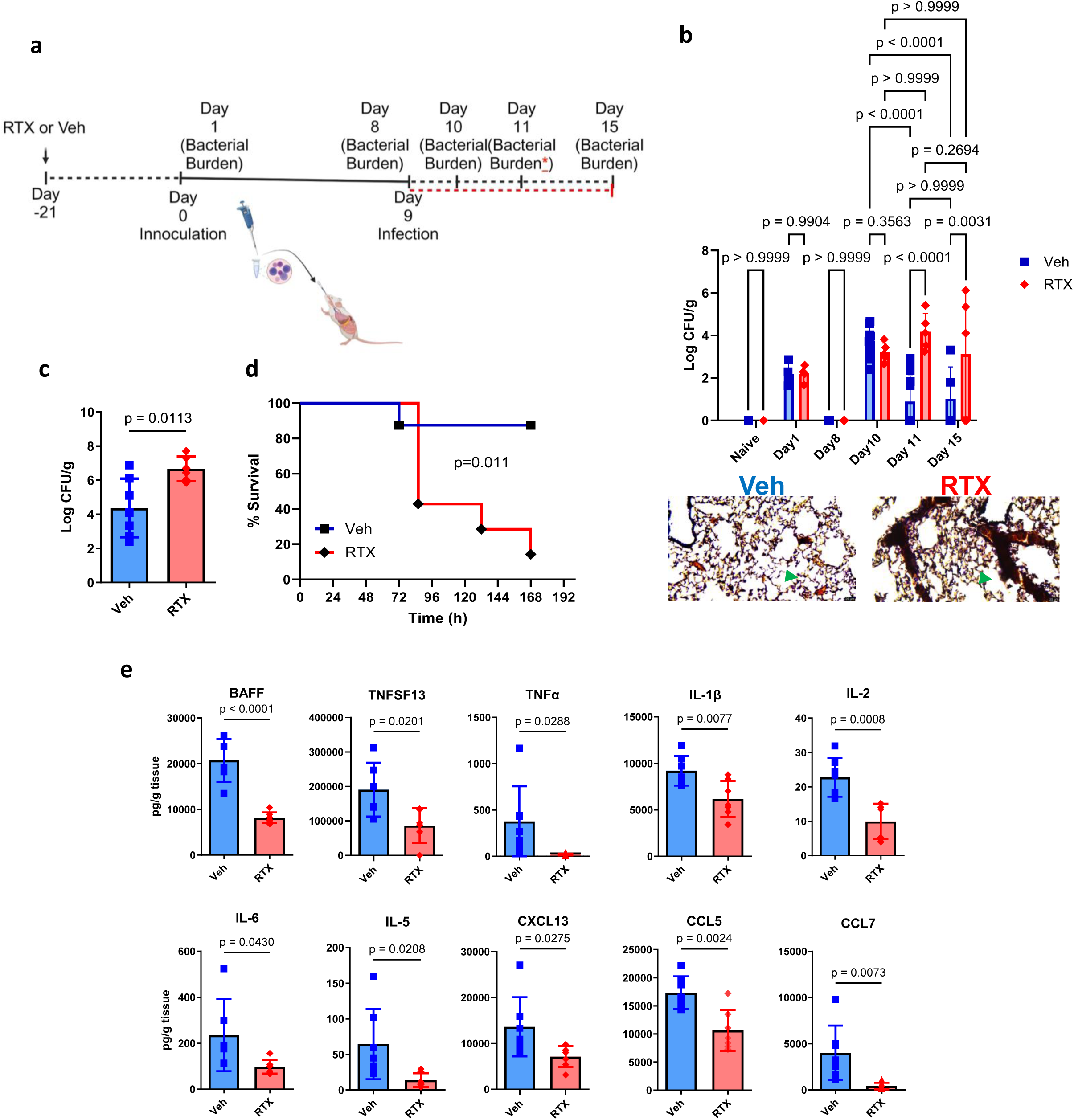
Sensory neurons are required for successful clearance of *S. pneumoniae* following pre-exposure and infection. **a**) Model of *S. pneumoniae* pre-exposure and infection. Mice received escalating doses of resiniferatoxin (RTX) or Vehicle (Veh) starting 21 days prior to inoculation. 10^4^ CFU of *S. pneumoniae (Serotype 19F ATCC49619)* in 50µl PBS was delivered on day 0 to expose mice, then 10^4^ CFU in 50µl PBS was delivered on Day 9. Bacterial burden was enumerated on Day 10, 11 and 15. Red asterisk and dotted line denote the separate set of experiments assessing cross-protection between serotypes: 10^4^ CFU of *S. pneumoniae (Serotype 19F ATCC49619)* was delivered on day 0 to expose mice, then 10^6^ CFU *(Serotype 3, ATCC6303)* in 50µl PBS was delivered on Day 9, survival was assessed as was bacterial burden on Day 11. **b)** Bacterial recovery from lungs in untreated sensory neuron intact mice (Naïve), untreated sensory neuron ablated mice (Naive RTX), pre-exposed and infected sensory neuron intact (Veh) or pre-exposed and infected sensory neuron ablated mice (RTX). Naïve n=6, Naïve RTX: n=5, Veh: n=5-13, RTX: n=5-7. Two-way ANOVA with Newman-Keuls post hoc test. Data were pooled from three independent experiments. Gram stain shows *S. pneumoniae* infection (purple and green arrows point to example of gram-positive space) was greater in RTX compared to Veh. Scale bar=20µm. **c)** Veh mice had reduced bacterial burden in response to primary inoculation with serotype 19F and infection with serotype 3 (Veh n=7, RTX n=6). Data were compared with two-sided t-test. **d**) 90% of Veh mice survived infection with serotype 3 compared to 20% of RTX mice after inoculation with 19F (Veh n=8, RTX n=7). Data were compared with Log-rank (Mantel-Cox) test. **e)** Quantification of B cell activating factor (BAFF), APRIL13 (TNFSF13), TNFα, IL-1β, IL-2, IL-6, IL-5, CXCL13, CCL5, CCL7 in sensory neuron intact (Veh n=7) and sensory neuron ablated (RTX, n=6) mice 16h following the final 10^8^ CFU dose of *19F*. Samples were analyzed with Luminex bead assay. IL-6 was re-run with ELISA. Two-sided t-test. Data were pooled from 2 independent experiments.

Next, we determined whether nociceptors regulated pro-inflammatory-cytokine production before lung infection clearance. Post-infection, RTX-treated mice showed reduced inflammatory cytokine levels, including IL-1β, IL-6, and TNFα (**Figure 1e, Supplementary** Figure 4).

Furthermore, cytokines involved in B-cell stimulation (BAFF, TNFSF13, IL-2, and IL-5) and chemokines involved in B-cell attraction (CXCL13, CCL5, and CCL7) were reduced in RTX mice compared to control mice (**Figure 1e, Supplementary** Figure 4).

### TRPV1+ neurons regulate B-cells

Given that cytokines and chemokines for B-cell recruitment (**Figure 1e**) were reduced, we assessed immunoglobulin levels in lung homogenates. Immunoglobulins were reduced in RTX mice compared to sensory neuron intact mice after 16hrs of infection (**Figure 2b**). *S. pneumoniae* serotype 19F specific IgG was also reduced in RTX as the IC50 calculated from lung homogenate dilutions was higher in RTX compared to Veh (**Figure 2b**). The reduction in immunoglobulins was directly due to a reduction in isotype-switched memory, resident memory B-cells, plasma cells, and plasmablasts (**Figure 2c-g, Supplementary** Figure 5-7). It is important to note that RTX did not affect B-cell populations immediately before pre-exposure or after first exposure (**Figure 2c-g**). The reduction in B-cell populations remained significant at 48h (**Figure 2c-g, Supplementary** Figure 6), in direct accord with an augmented bacterial burden in RTX mice (**Figure 1b,c**). As B-cells were diminished in RTX following infection, in accord with immunoglobulins, our data are consistent with a role for sensory neurons with adaptive immunity. In contrast, T-cells were unaffected by sensory neuron ablation (**Supplementary** Figure 8).

**Figure 2.**
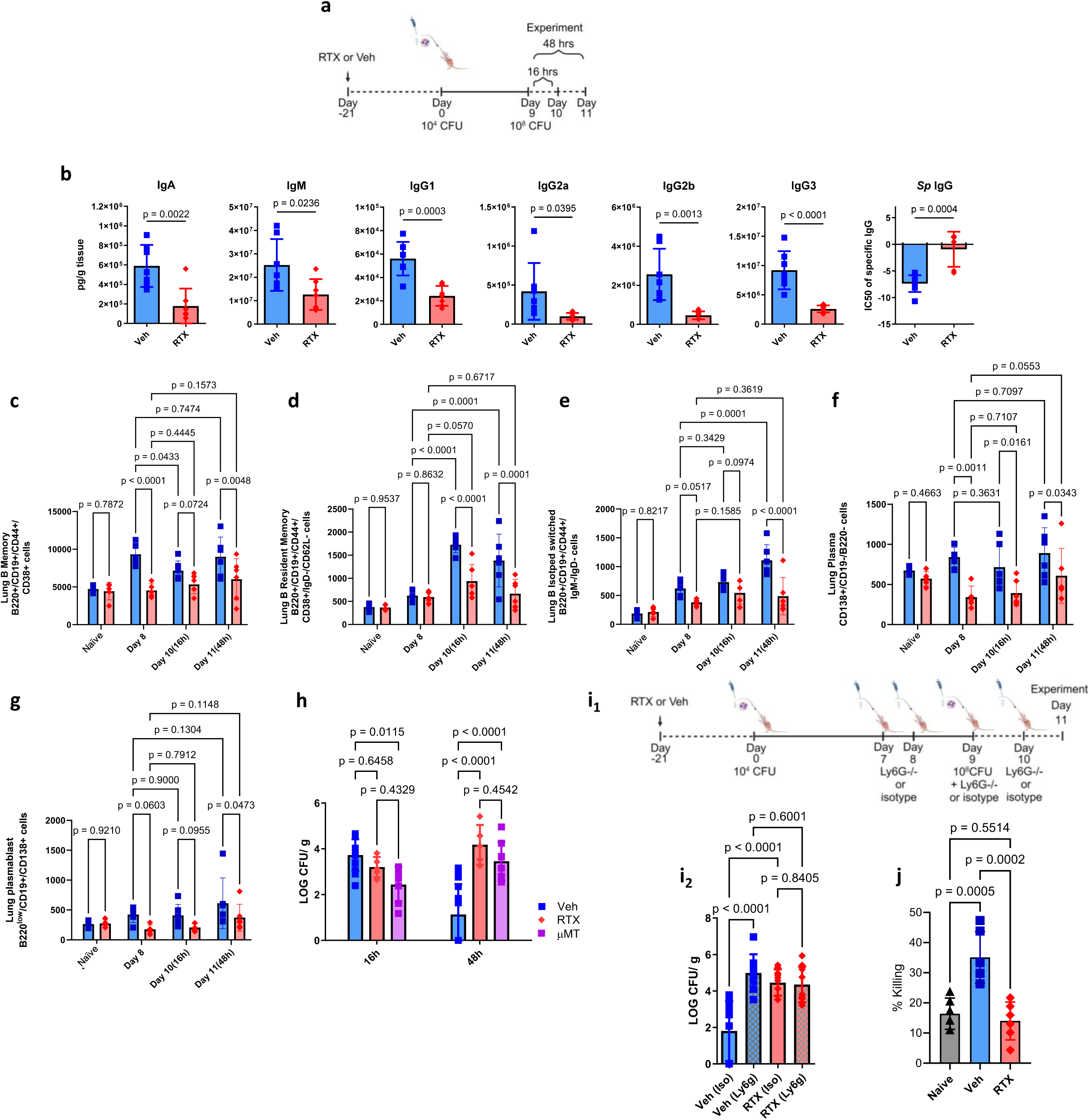
Sensory neuron ablation reduces, immunoglobulins, B-cells and disrupts neutrophil effector function after *S. pneumoniae*. **a**) In response to our pre-exposure and infection model (as in Figure 1a) **b**) IgA, IgM, IgG1, IgG2a, IgG2b, IgG3 were reduced with sensory neuron ablation 16h (Day 10) after final infection. Veh: n=7, RTX: n=7. Isotyping Luminex was used to analyze lung homogenates. Two-sided T-test. IgG specific to *S. pneumoniae* was tested with crude *Sp* extract. Serial dilutions were used to model the IC50 with a 4-point log model to determine differences between Veh (n=8) and RTX (n=6). Two-sided T-test. Data were collected from two independent experiments. **c**) B-memory B220+/CD19+/CD44+/CD38+ cells, **d**) B resident memory B220+/CD19+/CD44+/CD38+/IgD-/CD62l- cells, **e**) Isotype switched B220+/CD19+/IgM-/IgD- cells, **f**) Plasma CD138+ cells and, g) Plasmablast CD138+/B220-/CD19- cells were downregulated by sensory neuron ablation at 16 and 48h after infection. Only B memory and plasma cells were different between Veh and RTX on Day 8. Two-way ANOVA with Holm-Sidak test. Veh: n=5-7, RTX n=5-7. **h)** µMT mice had reduced burden at 16h in comparison to sensory neuron intact mice (Veh). Sensory neuron ablated (RTX) mice demonstrated a bacterial burden similar to mice with immature B cells (µMT) 48h after infection. Sensory neuron intact mice (Veh) reduced their bacterial burden at 48h. Veh n=13, RTX n=5, uMT infected n=7. Data from four independent experiments. Two-way ANOVA with Holm-Sidak tests. **i**) Ly6G A18 neutrophil depletion antibody or isotype control was delivered before, during and after the final infection dose of *S. pneumoniae.* Bacterial burden (Log CFU/ g lung mass) from lung homogenates was greater with neutrophil depletion in sensory neuron intact infected mice compared to isotype control 48h (Day 11) after infection. Sensory neuron depleted mice (RTX) did not further increase bacterial burden with neutrophil depletion demonstrating that B-cell neutrophil interaction was disrupted. One-way ANOVA with Holm-Sidak post-hoc test. Veh(Iso): n=12, Veh(Ly6g): n=8, RTX(iso): n=10, RTX(Ly6g):n=9. Data were pooled from two independent experiments. **j**) Naïve neutrophils were incubated with decomplemented serum harvested from naïve, Veh or RTX mice (pre-exposed and infected) and plated in S pneumoniae 19F coated wells (5000 CFU). Neutrophils were lysed 30 minutes later and plated on blood agar plates and CFU were enumerated 24h later. One-way ANOVA with Holm-Sidak post-hoc, n=6 per group.

To confirm that bacterial clearance in response to infection after pre-exposure relies on intact B-cells, we compared bacterial burden from our sensory neuron intact and depleted mice to mice lacking mature B-cells (µMT mice). Like RTX mice, µMT mice had a significantly increased bacterial burden at 48hrs after our recall model of *S. pneumoniae* (**Figure 2h**), Neutrophil depletion in sensory neuron intact mice significantly increased the bacterial burden to a level of RTX mice (**Figure 2i, Supplementary** Figure 9). Neutrophil depletion did not further augment bacterial burden in RTX mice, showing that the interaction between B-cells and neutrophils may be an important mechanism of bacterial clearance in our recall model (**Figure 2i, Supplementary** Figure 9). To confirm the B-cell-neutrophil axis, naïve neutrophils were incubated with decomplemented serum harvested from sensory neuron intact and RTX, which underwent our pre-exposure and infection model, as well as naïve mice. Neutrophils that were incubated with serum from sensory neuron intact mice had the highest level of bacterial killing (**Figure 2j**). Therefore, the main effect of sensory neurons in response to infection is to coordinate B-cell homing and immunoglobulin production to enhance neutrophil-mediated *S. pneumoniae* clearance.

### TRPV1+ neuron neuropeptide release of VIP stimulates B-cell immunoglobulin release

B- cells express select neuropeptide receptors^54^. Therefore, we reasoned that neurons would stimulate B-cells within their proximity. Using a sensory neuron reporter mouse (Vglutcre- tdTomato^50^), we showed that B220+ cells were localized within <20um of sensory neurons in the lungs 48h after the final infectious dose of *S. pneumoniae* (**Figure 3a**). In comparison, infected mice treated with RTX, showed a more dispersed distribution of B220+ cells, extending further from sensory neurons. Of note, whereas RTX depletes TRPV1 positive neurons, sensory neurons have a wide array of genetic composition^50,55^, and therefore, some nerves remain following TRPV1 depletion (**Figure 3a**). Thus, it was likely that sensory neuropeptides released by TRPV1+ neurons were suppressed with RTX. Therefore, we measured known sensory neurotransmitters with noted immunological effects (CGRP^2^, Substance P^10,11,56,56–58^, VIP^8,40,59,60^, NPY^35,42,61^) in lung homogenates. We show pre-exposure and infection increased NPY and VIP (**Figure 3b, c**). In comparison to sensory neuron intact mice, we demonstrate that RTX suppresses NPY, VIP, and CGRP (**Figure 3d, e**). The suppressed neuropeptide concentrations in the lung were associated with reduced cDNA expression in the vagal ganglion (**Supplementary** Figure 1). The neuropeptide concentrations measured from lungs are inversely correlated with bacterial burden and positively correlated with immunoglobulins (**Supplementary** Figure 10). Therefore, we hypothesized that NPY or VIP released from sensory neurons increase B-cell immunoglobulin release, to stimulate neutrophil-mediated bacterial clearance. Neuropeptides bind to specific receptors. We show that VIP1R and NPY1R are both present on B-cells and plasma cells, which demonstrates the likelihood of stimulation by sensory neuropeptides (**Supplementary** Figure 11).

**Figure 3.**
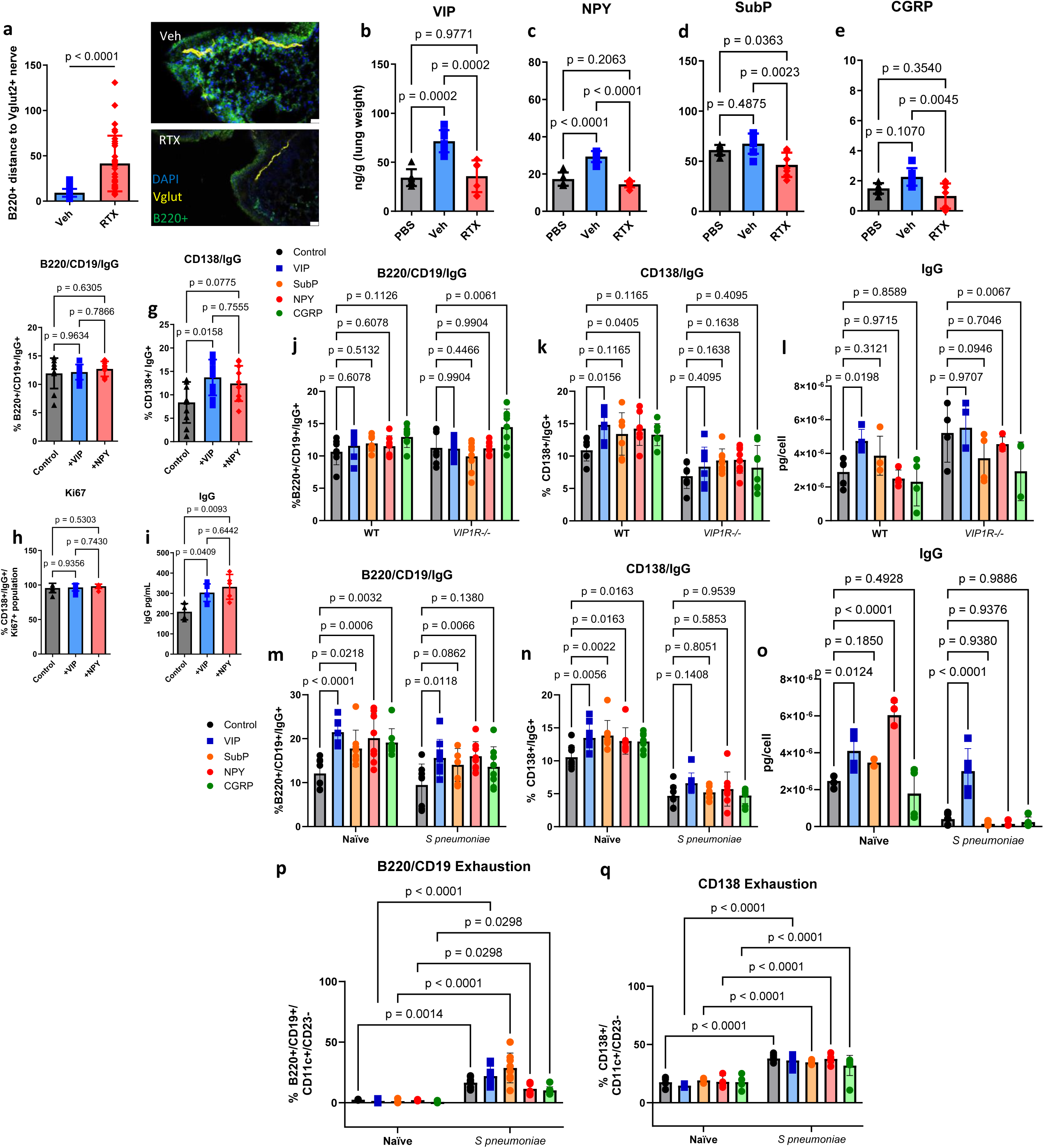
Vasoactive intestinal peptide attracts B-cells and regulates immunoglobulin production and release. **a**) B220+ cells cluster around sensory nerves from lungs of sensory neuron intact mice (Veh). In RTX sensory neuron depleted mice, B220+ cells are further dispersed. 30 cells (10 cells per sample) from n=3 mice for Veh & RTX. Two-sided t-test. **b-e)** Prominent sensory neuron neuropeptides were increased in lung homogenates 16 hours after pre-exposure and infection with *S. pneumoniae*. **b**) VIP was increased with infection and suppressed with sensory neuron depletion. **c**) NPY was increased with infection and suppressed with sensory neuron depletion. **d**) Substance P did not significantly increase following infection with *S pneumoniae.* **e**) CGRP did not significantly increase following infection with *S pneumoniae.* Lung homogenates analyzed with ELISA. PBS: n=6, Veh: n=7, RTX: n=7. One-way ANOVA with Tukey’s post hoc test. Data from 2 independent experiments. **f-i)** B cells were isolated from spleens of naïve mice and cultured in media with IL4+LPS and supplemented with VIP or NPY. Cells and media were sampled after 96 hours of incubation. **f**) VIP did not increase IgG bound to B220/CD19+ cells. **g**) VIP significantly increased bound IgG in CD138+ plasma cells. **h**) Proliferation as assessed by intracellular Ki67 staining was not affected by neuropeptides (from CD138+ gate). **i**) Secreted IgG was increased by VIP and NPY. n=5-10. One-way ANOVA and Tukey’s post-hoc test. Data from 2 independent experiments. **J-l**) B-cells were isolated from the spleens of VIP1 receptor knockout (VIP1R-/-) and WT littermates and cultured for 96h. **j**) B220/CD19+ cells were not significantly affected by VIP1R deletion. **k**) VIP1R-/- CD138+ cells did not significantly upregulate IgG in response to VIP and other neuropeptides. **l**) IgG released into the media was not increased with VIP or other peptides with VIP1R genetic deletion; VIP increased WT B-cell IgG release. Two-way ANOVA with Holm- Sidak post-hoc test. N=4-7. **m-p**) B-cells were isolated from Lungs of naïve or *S. pneumoniae* pre-exposed and infected mice. **m, n**) VIP increased bound IgG in B220+/CD19+ and CD138+ cells. This effect was greater in naïve cells. **o**) released IgG was increased with VIP in both naïve and pre-exposed and infected cells. **p,q**) Pre-exposed and infected cells show higher markers of B cell exhaustion after culture. Two-way ANOVA with Holm-Sidak post-hoc. Naïve n=4-7, S pneumoniae n=4-7.

We isolated splenic B-cells and cultured them with NPY or VIP. VIP stimulation increased IgG secretion (**Figure 3i**) and surface-bound levels in CD138+ plasma cells (**Figure 3f, g, Supplementary** Figure 12). However, neurotransmitter stimulation did not affect the proliferation of B-cells and plasma cells, assessed by Ki67+ intracellular staining (**Figure 3h**). Given the high expression of VIP1R on B-cells, we tested the effects of VIP stimulation in WT and VIP1R-/- isolated B-cells. In VIP1R-/- cultured B-cells, VIP and other neuropeptides did not increase bound and released IgG, demonstrating that VIP1R is the main receptor responding to VIP stimulation (**Figure 3j-l**). We then isolated B-cells from lungs of naïve, and pre-exposed and infected mice. VIP again increased bound and released IgG (**Figure 3m-o**). However, we noted that pre-exposed and infected mice had a suppressed response to all conditions. Therefore, we tested whether this suppression was due to B-cell exhaustion. B-cells harvested from pre-exposed and infected mice, which were then re-stimulated in culture for 96h, showed an augmentation of CD11c and reduction of CD23, to a greater extent than lung B-cells harvested and cultured from naïve mice (**Figure 3p, q**), indicative of B-cell exhaustion^62–64^.

We supplemented sensory neuron intact and RTX sensory-depleted mice with VIP to test a functional outcome of VIP release in response to *S. pneumoniae* (**Figure 4a, Supplementary** Figure 13). VIP supplementation rescued RTX mice by suppressing their bacterial burden at 48h post-infection (**Figure 4n**). Furthermore, B memory, plasma cells, plasmablasts, and lung IgG were increased with VIP supplementation in RTX mice compared to RTX mice supplemented with PBS (**Figure 4b-g**). Finally, *VIP1R*^−/−^ had an altered accumulation of select B-cell populations (**Figure 4h-k**), reduced immunoglobulins and cytokines (**Figure 4m, Supplementary** Figure 14), and increased bacterial burden in comparison to WT littermates in response to pre-exposure and infection (**Figure 4o**). In addition, when neutrophils were ablated with a neutralizing antibody, *VIP1R*^−/−^ mice did not further increase the bacterial burden in their lungs following pre-exposure and infection (**Figure 4p**). These data show that VIP stimulation of B-cells through the VIP1 receptor increases B-cell neutrophil interaction through an increased release of immunoglobulins.

**Figure 4.**
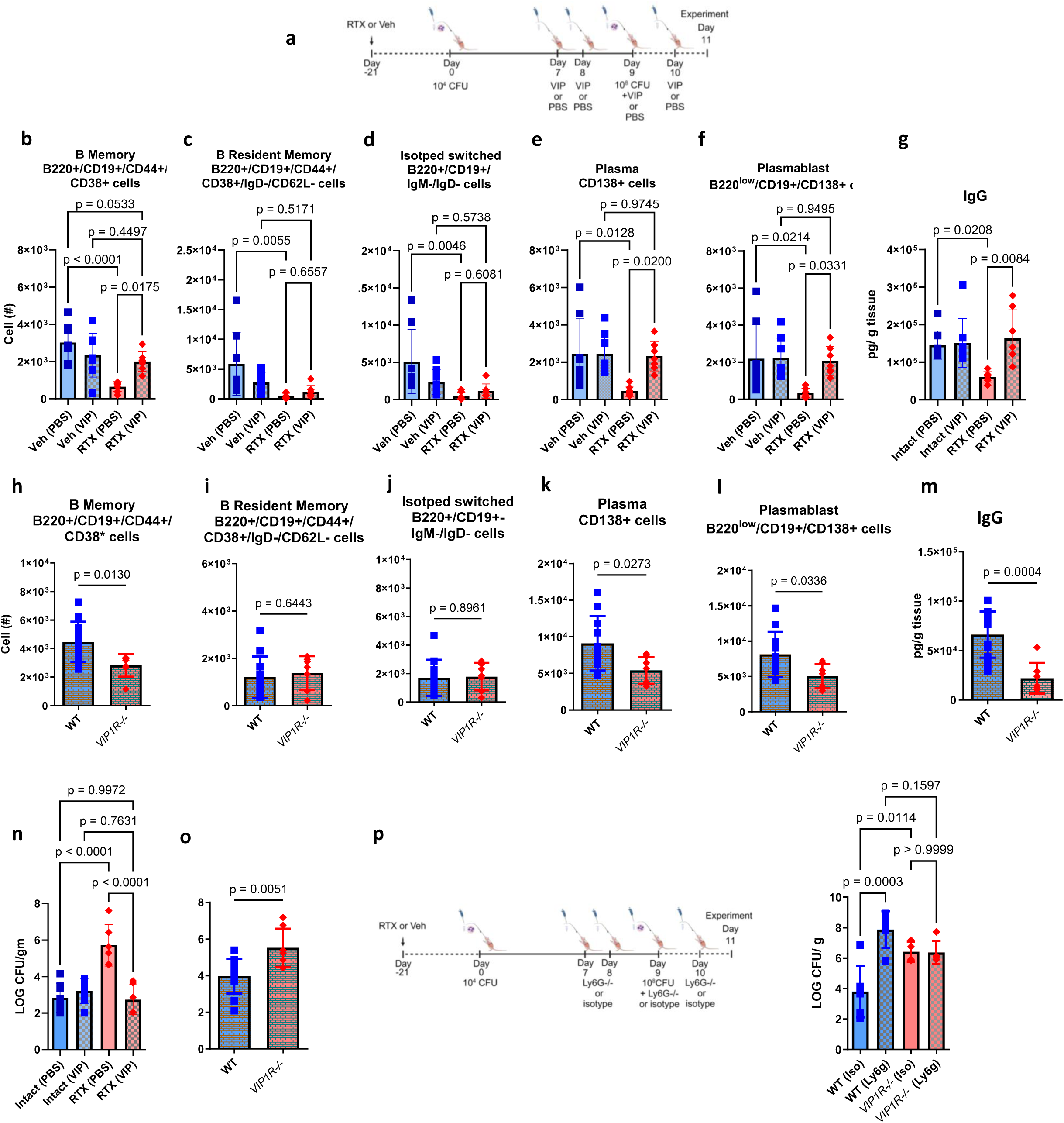
Vasoactive intestinal peptide supplementation improves bacterial clearance, memory B cells, plasma cells and IgG in response to pre-exposure and infection with *S. pneumoniae*. **a**) Vasoactive intestinal peptide (10ng/µl) or vehicle (PBS) was delivered as per the schematic in addition to the model of *S. pneumoniae* pre-exposure and infection with serotype 19F. Cells were gated on Live/CD45+ cells and total populations assessed with counting beads. **b**) B memory cells, **c**) B resident memory, **d**) Isotype switched, **e**) Plasma cells, **f**) plasmablasts were reduced with sensory neuron ablation. Supplementation with VIP increased B-cell populations. VIP supplementation did not further increase B-cell populations in sensory neuron intact mice. **g**) IgG was reduced with RTX as previous but increased with the supplementation of VIP to RTX mice. **h**) VIP 1 receptor knockout (VIP1R-/-) mice had reduced B memory cells compared WT mice, **i**) B resident memory and, **j**) Isotype switched B-cells were similar in VIP1R-/- and WT mice, **k**) Plasma cells, and l) plasmablasts were reduced in VIP1R-/- vs WT mice. Two-sided T-test. WT n=5, VIP1R-/- n=5. **m**) VIP1R-/- mice had reduced IgG following pre-exposure and infection compared to WT littermates. **n**) VIP supplementation suppressed bacterial burden in RTX mice. No effect of VIP supplementation was observed in Veh mice. **o**) VIP1R-/- mice have increased bacterial burden following pre-exposure and infection with *S. pneumoniae* compared to WT littermates. Veh (PBS): n=8, Veh (VIP): n=7-8, RTX(PBS): n=6-7, RTX(VIP): n=6-7. One-way ANOVA with Holm sidak post hoc test. Data from 3 independent experiments. WT n=8-11, VIP1R-/- n=5-7. Two-sided t-test. Data from 3 independent experiments. **p**) Ly6g neutralizing antibody or isotype was given to WT or VIP1R-/- mice. Bacterial burden (Log CFU/ g tissue mass) from lung homogenates was greater with neutrophil depletion in WT mice compared to isotype control. VIP1R-/- mice did not further increase bacterial burden with neutrophil depletion demonstrating that VIP significantly stimulates B-cells to increase neutrophil-mediated bacterial clearance. One-way ANOVA with Holm-Sidak post-hoc test. WT(Iso): n=6, WT(Ly6g): n=5, VIP1R-/-(Iso): n=4, VIP1R-/- (Ly6g):n=5. Data were pooled from two independent experiments.

In order to rule out non-neuronal sources of VIP, we stained for VIP in T-cells, macrophages and neutrophils which others have identified as potential non-neuronal sources of VIP^65–67^. RTX did not suppress VIP expression in leukocytes (**Supplementary** Figure 15). Neuroendocrine cells within the lungs also have the potential to release neuropeptides^68^. However, VIP fluorescence was not diminished with RTX in the epithelium where pulmonary neuroendocrine cells reside (**Supplementary** Figure 15). It is possible that these alternate sources of neuropeptides may contribute to B-cell stimulation. However, we note that in conjunction with the reduction of VIP in RTX lung homogenates (which selectively ablate neurons, **Figure 3b**), the suppressed expression of VIP transcripts in vagal ganglia in RTX compared to vehicle mice (**Supplementary** Figure 1) and a maintained amount of VIP-positive lymphocytes and neuroendocrine cells, the predominant source of neuropeptides in response to *S. pneumoniae* is likely lung TRPV1+ sensory neurons.

### Neural stimulation of immunoglobulins exacerbates asthma features

Given that immunoglobulins and humoral immunity are important mechanisms for bacterial clearance, we then asked whether an overproduction of immunoglobulins as a result of neural stimulation could result in immune dysfunction. We hypothesized that this would be neuropeptide driven but, likely mediated by an alternate peptide as previous data supports^10,58^. *A. alternata* causes a swift and distinct rise of IgE in response to four inoculations (**Figure 5a**) as demonstrated previously^69^. *A. alternata*-induced asthma increased airway hyper-responsiveness and goblet cell metaplasia, which are reduced in sensory neuron depleted mice (**Supplementary** Figure 16b-e) where airway hyper-responsiveness was correlated with IgE (**Supplementary** Figure 16c). Specific B-cell recruitment of cytokines is suppressed with RTX sensory neuron ablation (**Supplementary** Figure 17). RTX also suppressed B-cells, immunoglobulins, and mast cells in asthmatic mice (**Figure 5c-h, Supplementary** Figure 18). *A. alternata* specific IgE was also reduced in RTX mice as the IC50 calculated from lung homogenate dilutions was higher in RTX compared to Veh mice(**Figure 5b**). NPY and Substance P were increased in the lungs during *A. alternata*-induced asthma, in comparison to naïve mice, while decreased with RTX (**Supplementary** Figure 19a-d), VIP was not increased in response to *A. alternata*-induced lung inflammation (**Supplementary** Figure 19d). Substance P receptors (Neurokinin 1 and 2) were not present in our isolated naïve B-cells (**Supplementary** Figure 11) but CDNA was detectable in lung B-cells. Substance P also binds to mas-related g protein-coupled receptors^70^, where cDNA and protein were present on all B-cells (**Supplementary** Figure 11). NPY and Substance P increased bound and released IgE in naïve splenic B-cells (**Supplementary Figure12, 19e-h**). To test whether substance P could stimulate B-cells, we supplemented substance P into *A. alternata* mice (**Figure 6a, Supplementary** Figure 20). We show that substance P supplementation increased B-cell resident populations in the lungs of RTX sensory neuron-depleted mice treated with *A. alternata* (**Figure 6b-f**). Further, IgE was increased when *A. alternata* RTX mice were supplemented with substance P (**Figure 6l**). This was in line with the removal of substance P, as substance P knockout mice (unable to produce substance P/ have receptors) showed reduced levels of IgE (**Figure 6m**), lung-resident B-cell populations (**Figure 6g-k**), as well as cytokines and other immunoglobulins (**Supplementary** Figure 21) in response to *A. alternata*-induced lung inflammation compared to WT mice with *A. alternata- induced* lung inflammation. To account for alternate sources of substance P, we stained for substance P in T-cells, eosinophils, and mast cells, which could be sources of substance P^65–67^. RTX did not suppress substance P expression in leukocytes (**Supplementary** Figure 22). We note that substance P fluorescence was not diminished with RTX in the epithelium where neuroendocrine cells are located (**Supplementary** Figure 22). In conjunction with the reduction of substance P in RTX lung homogenates (**Supplementary** Figure 19), the suppressed expression of substance P transcripts in vagal ganglia in RTX compared to vehicle mice (**Supplementary** Figure 1) and a maintained amount of substance P positive leukocytes and epithelium, we believe the predominant source of substance P in response to *A. alternata* are lung sensory nerves. It is still possible that these alternate sources of neuropeptides, including leukocytes and neuroendocrine cells, could contribute to B-cell stimulation. In summary, these data demonstrate that substance P exacerbates asthmatic features in mice by increasing B-cell production of IgE.

**Figure 5.**
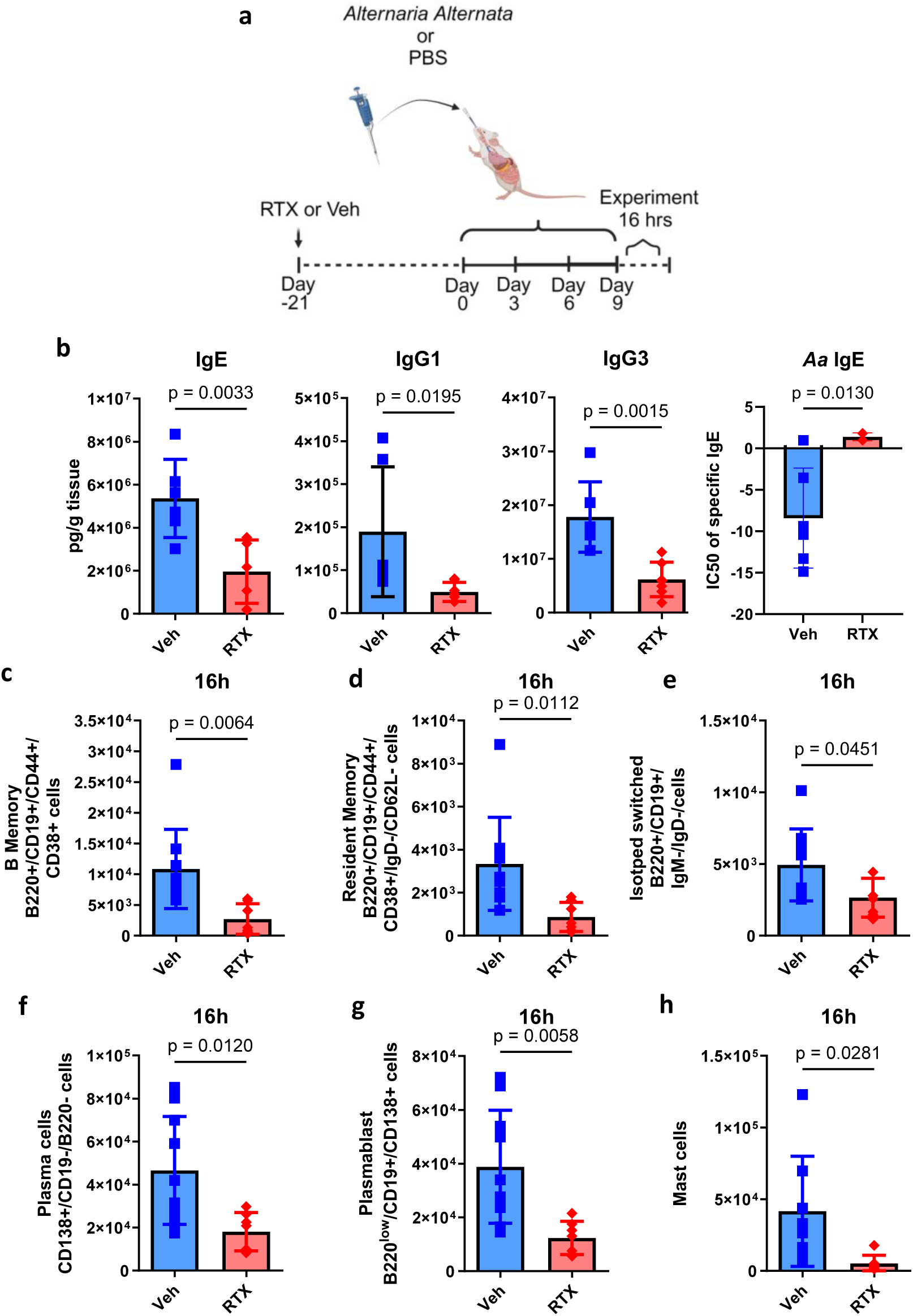
Sensory neuron ablation reduces B-lymphocytes, immunoglobulins and, mast cells in *A. alternata* treated mice. **a**) *A. alternata* induced asthma was elicited as per Cavagnero et al. 25µg of *A. alternata* extract in 50µl PBS was delivered as per the schematic. Experiments took place 16hrs later. **b)** IgE, IgG1, IgG3 (n=6, RTX: n=7) were reduced with sensory neuron ablation after final *A. alternata dose*. Isotyping Luminex was used to analyze lung homogenates. *A. alternata* extract with lung homogenates and IC50 modeling was used to determine IgE specific to *A. alternata* (Veh n=6, RTX n=4) Two-sided t-test. Data were collected from two independent experiments. **c**) B-Memory B220+/CD19+/CD44+/CD38+ cells, **d**) B resident memory B220+/CD19+/CD44+/CD38+/IgD-/CD62l- cells, **e**) Isotype switched B220+/CD19+/IgM-/IgD- cells, **f**) Plasma CD138+ cells and, **g**) Plasmablasts CD138+/B220-/CD19- cells and **h**) mast cells CD117+ were downregulated by sensory neuron ablation 16h after *A. alternata* asthma induction. Cells were gated on live/CD45+ cells and counting bead calculation determined total cell population. Red gate is the measured cell population. Veh: n=10, RTX: n=7. Two-sided t-test. Data from 3 independent experiments.

**Figure 6.**
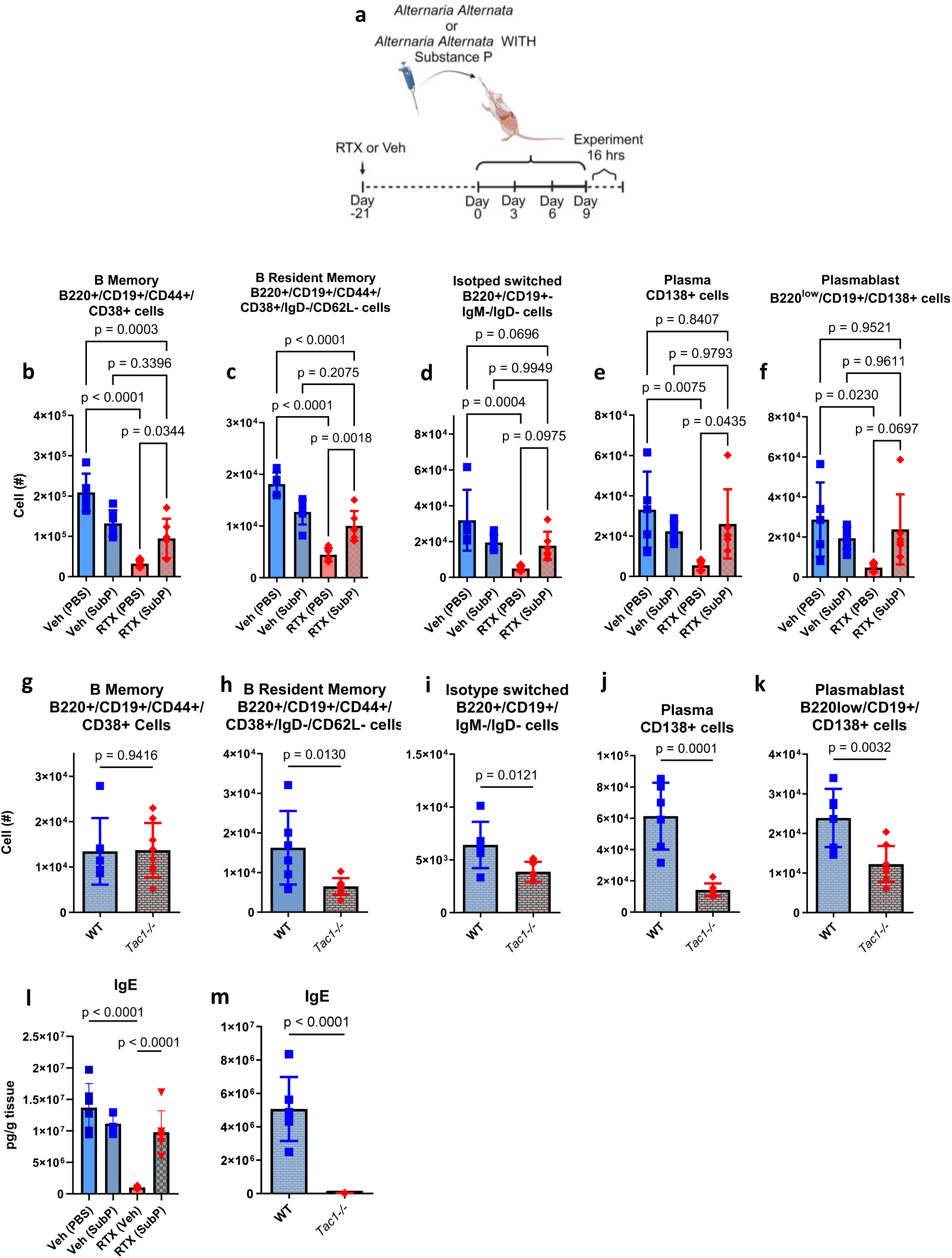
Substance P increases B-cells and IgE in mice treated with *A. alternata*. **a**) Substance P (100ng/µl) or vehicle (PBS) was delivered in conjunction with *A. alternata* extract as per the schematic, in accordance with our *A. alternata* model of asthma. Cells were gated on Live/CD45+ cells and total populations assessed with counting beads. **b**) B memory cells, **c**) B resident memory, **d**) Isotype switched, **e**) Plasma cells, **f**) plasmablasts were reduced with sensory neuron ablation. Supplementation with Substance P increased B-cell populations.Substance P supplementation did not further increase B-cell populations in sensory neuron intact mice. Veh (PBS): n=5, Veh (Sub P): n=6, RTX(PBS): n=6, RTX(SubP): n=6. One-way ANOVA with Holm sidak post hoc test. Data from 3 independent experiments. Tac1-/- (lack Tac1 gene, unable to produce substance P, have receptors) had similar B memory (**g**), but reduced B Resident Memory (**h**), Isotype switched (**i**), plasma cells (**j**), and plasmablast cell (**k**) compared to WT mice receiving *A. alternata* induction of asthma. WT n=6, Tac1-/- n=8, Student’s unpaired t-test. Data from 2 independent experiments. **l**) Supplementation with substance P increased IgE in *A. alternata* mice treated with RTX to a level consistent with Veh. Substance P did not further increase IgE in Veh mice (All groups n=6). One-way ANOVA with Holm sidak post hoc test. Data from 2 independent experiments. **m**) Substance P knockout mice (Tac-/- n=7) had reduced IgE compared to WT *A. alternata* mice (n=8). Two-sided t-test. Data from 2 independent experiments.

## Discussion

Models of repeated immune stimulation show that sensory neurons may stimulate humoral immunity^10^. In contrast, sensory neurons appear to suppress innate responses to select single-dose models of bacterial and fungal infection^1–7^. Recent data show that sensory neurons modulate B-cell populations in the lungs^10^ and contribute to B-cell maturation in the mucosa^39,43^. Select neuropeptides, released by sensory neurons, can induce IgE, IgG, and IgA release *in vitro*^34–38^. Therefore, we reasoned that sensory neurons could play critical roles in humoral immunity^10,71–73^. The differential roles of neurons with innate and adaptive immunity may explain why sensory neurons can produce divergent results depending on the underlying immune stimulus, which is likely dependent on the timing and type of neuropeptide released.

Sensory neurons are able to sense pathogenic infiltration directly through PAMPs or indirectly by binding cytokines and proteases^21–23^. Upon stimulation, sensory neurons release neuropeptides directly into the immediate vicinity^21–23^. The type and amount of neuropeptide released is likely dependent on the neuron type stimulated, which would dictate how it senses the environment and which neuropeptides would be released^55,74^. Depletion of sensory neurons reduced B-cell populations, IgG, and neutrophil activity, which subsequently augmented bacterial burden in our recall model. This humoral regulation was due to the release of VIP, likely from sensory neurons and the resulting stimulation of B-cells and plasma cells via the VIP1 receptor. We demonstrate that B-cells and plasma cells have VIP1 receptors, which stimulate PKCs that are critical to the development and release of immunoglobulins^75,76^. These data are confirmed with VIP1 receptor knockout mice, highlighting the essential role of VIP1 receptor signaling in the proper functioning and immunoglobulin-secreting capability of B-cells and plasma cells. While RTX treatment suppresses neuropeptide gene transcripts in sensory neurons and reduces neuropeptide levels in lung homogenates, sensory neuropeptides may still activate the release of neuropeptides from other sources, such as lymphocytes and neuroendocrine cells. This potential alternative source of neuropeptide release means that our current analysis might underestimate the overall stimulation of B-cells by neuropeptides, further underscoring the importance of neuropeptide stimulation of B-cells.

Our results are consistent with early *in vitro* data which show that VIP can stimulate B-cell immunoglobulin release^34–38^. In the context of bacterial lung infection, our results contrast with the finding that sensory neurons suppress immunity in response to acute infection^2^. Specifically, sensory neuron release of CGRP suppressed bacterial clearance of acute pulmonary *Staphylococcus aureus infection* by dampening γδT cell-neutrophil stimulation^2^. When we used a single dose of *S. pneumoniae*, we saw that sensory neuron depletion had a similar bacterial burden and IgG as sensory neuron intact mice 48 hours following the single challenge as well as survival in response to a lethal strain. In response to pre-exposure and infection, CGRP did not significantly increase. We suggest that CGRP may be necessary for early innate neuroimmune interaction, which is superseded by VIP upon re-infection, or that CGRP may have an alternate role for B-cell maturation in germinal centers^77^. While humans are colonized with pneumococcal strains at birth, mouse models appear to require pre-exposure prior to infection in order to engage humoral memory and mimic the human condition^30,31,51,52^. Therefore, our model of pre-exposure and infection may provide data that is more applicable to the translation to clinical practice.

B-cells can become dysregulated and exacerbate diseases. In the context of allergen-induced lung inflammation, we show that sensory neurons become hypersensitized and stimulate B-cells. This is in direct accord with many investigations demonstrating that hypersensitization of sensory neurons leads to augmented cytokine release and subsequent eosinophil trafficking by T-cells or innate resident lymphocytes^8,9,28^. Our study extends these findings by showing that sensory neuron depletion reduces B-cell-attracting cytokines and, as a result, reduces B-cell recruitment. Mast cells were also reduced in sensory neuron-depleted mice. It is likely that the combined reduction in B-cells and IgE release also reduced mast cell stimulation. Conversely, mast cells also harbor neuropeptide receptors^78^, and sensory neuron ablation could directly reduce mast cell stimulation.

Substance P released by sensory neurons regulates B-cell stimulation in our asthma model. We and others show the ability of substance P to induce class switch recombination^10^. We demonstrate that mas-related gene protein-coupled receptors that are stimulated by substance P^11^ are ubiquitously expressed by B-cells with tachykinin receptor transcripts detectable in pulmonary B-cells. Our data significantly extend previous data by showing that the chemical ablation of neurons reduced B-cell populations and IgE levels. Further, when substance P was supplemented in asthmatic sensory neuron-depleted mice, IgE was once again restored. Finally, genetic ablation of the ability to produce substance P completely prevented IgE production in response to *A. alternata.* Increased substance P production has been associated with an asthmatic phenotype. Specifically, increased substance P-expressing neurons in asthma^79^ augmented mucus secretion and goblet cell hyperplasia^80–82^, which contributes to overall airway hyperresponsiveness^8^, as we show. Tachykinins such as substance P, therefore, contribute to asthma induction via direct airway stimulation and through IgE release.

While the direct effects of neuropeptides on B-cells have been demonstrated, recent findings show that the lymph node innervation by sensory neurons is restricted to the macrophage-rich medulla, not the lymphocyte-rich cortex^71,77,83^. Therefore, sensory neurons or neural circuits are unlikely to recruit B-cells from lymph nodes or secondary lymphoid tissues. In line with this reasoning, we show no changes in B-cells in the spleens or bone marrow in response to sensory neuron depletion. Our data demonstrate that the effects of sensory neuropeptides are confined to the lungs and produce these effects locally. Lung tissue-resident B-cells establish themselves after primary infection or colonization^84,85^, and are poised to release antibodies^86,87^ upon antigen encounter with a challenge infection. Our data demonstrate that B-cells cluster around neurons in lungs and that the changes to B-cell numbers were only in the lungs. Further, CD62L negative B resident cells were prominent in our model and reduced with sensory neuron ablation, demonstrating that tissue-resident B-cells were employed upon infection after pre-exposure. In contrast to the lengthy time course required for germinal center-mediated B-cell maturation, our rapid antibody responses were consistent with tissue-resident extra-follicular B-cell antibody production^88,89^. We suggest that neuropeptides act to assist in the maturation and development of tissue-resident B-cells.

Our data provides a new role of sensory neuropeptides acting as B-cell stimulants and highlights the humoral influence of sensory neurons. This study demonstrates that sensory neurons are critical in regulating pulmonary humoral immunity and the outcomes of bacterial lung infections and asthma. Targeting neuro-immunological communication through VIP and substance P or other potentially uncovered molecular mechanisms may be an effective approach to enhance host protection or reduce allergic responses by influencing immunoglobulin production through accessory pathways.

## METHODS

### Mice

All animal experiments were approved by The Lundquist Institute at Harbor UCLA Institutional Animal Care and Use Committee protocol # 32183. Mice were housed in a specific-pathogen-free animal facility at The Lundquist Institute. C57BL/6J, Tac-/- (B6.Cg- Tac1tm1Bbm/J), and µMT (B6.129S2-Ighmtm1Cgn/J) mice were purchased from Jackson Laboratories. Vglut-TdTomato mice were provided by X. Sun (UCSD- crossed from Vglut2cre- and RosaTdTomato originally attained from JAX B6J.129S6(FVB)-Slc17a6tm2(cre)Lowl/MwarJ and B6.Cg-Gt(ROSA)26Sortm14(CAG-tdTomato)Hze/J). TRPV1-DTR mice were obtained from Dr. Isaac Chiu, with MTA provided by M Hoon (NIH). VIP1R-/- were kindly donated by Joseph Pisegna, Patrizia Germano, and James Waschek (UCLA, MGI: 177616). At the start of experiments, mice was 5-8 weeks old. Age-matched male and female mice were used for experiments.

### Streptococcus pneumoniae cultures

*A. S. pneumoniae* strain 49619 serotype 19F and strain 6303 serotype 3 were purchased from ATCC. Cultures were originally grown on Tryptic Soy Agar (BD Biosciences 236950) supplemented with 5% defibrinated sheep’s blood (Remel R54012) and cultured for 24h, then re-cultured for an additional 24h in 5% CO_2_ at 37°C. Colonies were then picked for shape and hemolytic activity and cultured in Todd Hewitt Broth supplemented with 2% Yeast Extract (Fisher 50489152) or Brain Heart infusion media (BD DF0037178) in 5% CO_2_ at 37°C. Cultures were adjusted to OD 600 1.0 and resuspended in PBS prior to inoculation.

### Infection model

Colonies were checked for OD, then spun at 4000 rpm, 10 min, and washed in PBS. On day 0, mice were infected with 10^4^ CFU of Serotype 19F in 50 µl of PBS, intranasally. Then on day 9, mice were infected with 10^8^ CFU of serotype 19F in 50 µl of PBS intranasally. Bacterial burden were assessed in naïve mice (prior to any inoculation), immediately following the first inoculum (Day 1), prior to the second infectious dose (Day 8), 16 hours after the infectious dose (Day 10), 48 hours after the infectious dose (Day 11) and 6 days after the final infectious dose (Day 15). In a separate set of experiments serotype cross-protection was tested. Mice were infected with 10^4^ CFU of serotype 19F in 50 µl of PBS, intranasally. Then on day 9, mice were infected with 10^6^ CFU of serotype 3 in 50 µl of PBS intranasally., Survival and bacterial burden (at 48h post-infection) were assessed. Control animals were intranasally infused with 50μl PBS only.

### Allergy model

*A. alternata* extract was purchased from CiteQ Laboratories (09.01.26) and dissolved to yield a final concentration of 25ug/ml. Then mice were inoculated intranasally with 50ul on days 0, 3, 6, and 9 where experiments took place 16 or 48 hours later^90^. All control mice received PBS in the same volume intranasally.

### Sensory neural ablation

For chemical ablation of TRPV1+ neurons, mice 5 weeks of age were treated with RTX (Adipogen 502053716) as previously described^2,4,7^. Mice were injected subcutaneously in the dorsal flank on consecutive days with three increasing doses of RTX (30, 70, and 100 μg/kg) dissolved in 2% DMSO with 0.15% Tween 80 in PBS. Control mice received vehicle.

For targeted TRPV1+ neuron depletion, we injected 20 ng DT in 200 nl PBS containing 0.05% Retrobeads (LumaFluor INC) into nodose/jugular/petrosal VG with a nanoinjector (Drummond Scientific Company)^91^. Mice were anesthetized with 87.5mg/kg ketamine and 12.5mg/kg xylazine. The vagal ganglion was exposed after a midline incision in the neck (∼1.5 cm in length). DT was gently injected at a rate of 5nl/s. Once completed, the injection needle remained in the ganglia for 2-3 minutes in order to reduce the spillover of DT. This process was repeated for the vagal ganglion on the other side of the body.

### Bacterial burden determination

Lungs were weighed and then homogenized in 2ml sterile PBS with a tissue homogenizer. Homogenates were serially diluted on tryptic soy agar plates with 5% sheep’s blood and 10ug/ml neomycin (Fisher Biochemicals BP2669-5). The bacterial CFU were enumerated after overnight incubation at 37°C and 5% CO2.

### Neurotransmitter dosing

Vasoactive intestinal peptide (200ng in 50 μl PBS) was delivered intranasally at 48, 24, and 0h before and 24h after (for 48h harvest only) final infection with 10^8^ CFU of *S. pneumoniae*. Control mice received PBS at the same time points. When mice received bacteria on the same day, VIP was combined with the bacterial inoculum.

Substance P (1000ng in 50uL PBS) was delivered intranasally on Days 5,6,7,8,9 of the *A. alternata* model. On days where *A. alternata* was delivered, Substance P was mixed with *A. alternata*.

### Neutrophil depletion

For neutrophil depletion, we followed an established protocol^92^. Mice were injected i.p. with 150 μg of anti-A18 (clone 1A8, BioXCell BE-0075-1) (in 200 μl) per mouse 48, 24, 0h before lung infection and 24h after lung infection (48h analysis only). Control mice received 150 μg of rat IgG isotype control (2A3 BioXCell).

### Neutrophil bacteriocidal assay

Mouse neutrophils were purified as described previously^93^. In brief, bone marrow cells were flushed from femurs and tibias of 8 week old C57BL/6J mice using sterile RPMI 1640 medium supplemented with 10% FBS and 2 mM EDTA onto a 50 mL screw top Falcon tube fitted with a 100 μm filter. Mouse neutrophils were purified from bone marrow cells using negative magnetic bead selection (MoJo Sort 480057, BioLegend) according to the manufacturer’s instructions. Bone marrow-enriched neutrophils had >98% purity and>93% viability. Neutrophil killing of S*. pneumoniae* 19F was determined by CFU enumeration. *S. pneumoniae* 19F was incubated with 50% decomplemented (20 min; 65°C) serum in PBS from naïve, primary infected, and primary infected mice treated with RTX for 30 min on ice. Bacteria were washed with PBS, and 1x10^3^ serum-coated *S. pneumoniae* 19F were incubated with 1x10^5^ BM-neutrophils for 30 min. Neutrophils were lysed with 0.02% Triton X-100 in ice-cold water for 5 minutes, diluted, and the remaining bacterial cells were quantitatively cultured.

### Flexivent lung function

Mice anesthetized with ketamine/xylazine (87.5/12.5mg/kg), paralyzed with pancuronium bromide (MP Biomedicals ICN15605350), and subject to methacholine challenge with 200µl of doubling doses of methacholine (MP Biomedicals ICN19023110) dissolved in PBS for 20 breaths. Single and double compartment model Respiratory resistance was measured with the Flexivent system (SCIREQ) as previous^91^. Both absolute and percentage change values from PBS were calculated.

### Tissue collection

For flow cytometry, mice were euthanized by Isoflurane inhalation. The lungs were then dissected and flushed with PBS, coarse dissected, and incubated at 37°C for 45min in 1.75mg/ml collagenase IV (C4-22-1g Sigma) in PBS; then washed, macerated through a 21 gauge needle and filtered through a 70 µm mesh filter, then treated for flow cytometry. Spleens were macerated through a 70 µm mesh filter and then treated for flow cytometry. Bone marrow was flushed with RPMI media supplemented with 2mM EDTA and 10% FBS, filtered through a 70 µm mesh filter, then treated for flow cytometry.

For cytokine analysis, lungs were dried, weighed, then macerated in 2ml PBS supplemented with 25ul protease inhibitors (consisting of AEBSF HCl (100 mM), Aprotinin (80 μM), Bestatin (5 mM), E-64 (1.5 mM), Leupeptin (2 mM) and Pepstatin (1 mM) Tocris 5500), spun at 8000g 4 min 4C and then the supernatant was snap frozen and stored t -80°C until analysis.

### B-cell isolation

B-cells were isolated from spleens or lungs (below) using the EasySep Mouse B-cell negative selection kit. CD138+ cells (for qPCR, below) were isolated using the EasySep Mouse CD138 positive selection kit.

### Quantitative real-time PCR

RNA was isolated from tissues or cells using the inTron Easy spin Total RNA extraction Kit (Boca Scientific 17221), which was reverse transcribed to cDNA with a Tetro cDNA Synthesis Kit (Tetro biosystems NC1352749). Relative gene expression was determined by quantitative real-time PCR on a QuantStudio 3 System (Thermo Fisher Scientific) with TaqMan Fast Advanced Master Mix (Thermo Fisher Scientific 4444557) with the TaqMan probe sets (ThermoScientific Applied Biosystems). Expression values relative to HPRT were calculated by Δ ΔCTT method. All taqman probes utilized were purchased from Thermofisher Vip1R Mm00449214_m1; Vip2R Mm01238618_g1; Tac1R Mm00436892_m1; Tac2R Mm01175997_m1; NPY1R Mm00650798_g1; NPY2R Mm01956783_s1; NPY4R Mm00435894_s1; NPY5R Mm02620267_s1; NPY6R Mm00440546_s1; MRGPRA Mm01984314_s1; MRGPRB2 Mm01956240_s1; MRGPRG Mm01701870_s1; RAMP1 Mm00489796_m1; Tac1 Mm01166996_m1; Calca Mm00801463_g1; VIP Mm00660234_m1; NPY Mm01410146_m1; TRPV1 Mm01246300_m1; HPRT Mm03024075_m1).

### Cytokine, neurotransmitter, and immunoglobulin analysis

Cytokine and immunoglobulin levels in lung homogenates were measured through Luminex multiplex assay using R&D discovery or Thermo kits; analytes were measured with the Luminex 200 system. Enzyme- linked immunosorbent assay (ELISA) kits were used according to the manufacturer’s instructions (R&D systems, Raybiotech, or Thermofisher) and analyzed with a BioTek Synergy H1 plate analyzer. Luminex and ELISA kits are all commercially available and listed in the supplementary materials list.

### Specific immunoglobulin analysis

For specific IgG determination, crude S. pneumoniae (12.53µg/ml ATCC 49619) extract was incubated in 96 well plates overnight at 4°C, then blocked with goat serum for 1h at 4°C. Then log dilutions of lung homogenates (from groups above) were incubated at RT for 2 hours. Goat anti-mouse IgG HRP conjugated secondary antibody (Abcam ab205719, 1:1000) was then added and incubated for 2 hours RT. The ELISA was developed with 0.5% TMB solution (Fisher AAJ61325AP) and stopped with 2N H2SO4 (Fisher 828016). For specific IgE determination, *A. alternata* (100µg/ml CiteQ 09.01.26) was incubated as above and developed as above, except Goat anti-mouse IgE HRP secondary (Thermofisher PA1-84764, 1:1000) was used. OD450 for each dilution was used with a 4-point sigmoidal curve to determine IC50 for each sample in each group.

### Flow cytometry

Red blood cells were lysed with ACK lysing buffer (Gibco A10492-01), treated with Fc Block (Biolegend Trustain 101320), and resuspended in FACS buffer (HBSS-Gibco 10010-023 with 2% FBS Gibco. Incubations with antibody cocktails were conducted at 4°C for 60 min, and samples were subjected to two washes and resuspension in FACS buffer (HBSS +2% FBS Sigma F8192). For intracellular staining, cells were fixed/permeabilized with BD cytofix/cytoperm kit (554714), washed, and stained overnight at 4°C. Flow cytometry was conducted on a Symphony A5 flow cytometer (BD). Data were collected with BD DIVA software, and files were analyzed with FlowJo (Treestar, version 10.0.8r1). A live-cell stain (APC-Cy7, Invitrogen) was used to exclude dead cells. Gating strategies are provided in **Supplementary** Figure 5. Positive staining and gates for each fluorescent marker were defined by comparing full stain sets with fluorescence minus one (FMO) control stain sets. All antibodies were used at 1:100. Antibodies used: Biolegend-CD11c (Clone N418), Ly6g (Clone 1A8), CD117 (Clone 2B8), Siglec F (Clone 1RNM44N Invitrogen), F4/80 (Clone BM8), CD3 (BUV805), CD3 (Clone17As), CD11c (Clone N48), CD4 (Clone GK1.5), CD8 (Clone SK1), T-bet (Clone 4B10), GATA3 (Clone 16E10A23), B220 (Clone RA3-6B2), CD19 (Clone 6D5), CD62L (Clone MEL-14), CD44 (Clone IM7), CD138 (Clone 281-2), IgG1 (Clone RMG1-1), IgE (Clone RME-1) CD45 (Clone 30-F11), MRGPRX (Clone K125H4); BDbiosciences- IgM (Clone IL/41), IgD (Clone 11-26C.2a), RorγT (Clone Q31378; Invitrogen- CD38 (Clone 90), Ki67 (Clon SolA15); AlomoneLabs- VIP1R (Clone AB_2341081), Tac1 (polyclonal Proteintech), VIP (polyclonal Proteintech).

All in vivo cells were gated on leukocytes, singlets, live, CD45. Mast cells were CD117+/MHCII+. Netrophils were Ly6g+, Eosinophils were SiglecF+/CD11b+. T cells were divided into CD4+ or CD8+ fractions and intracellularly stained for Tbet (Th1), GATA3 (Th2) or RORgammaT (Gamma delta). B-cell lineage were B220+/CD19+. These were subdivided based on B Memory (CD44+/CD38+), B Resident Memory (CD44+/CD38+/IgD-/CD62L-), Isotype switched (IgM-/IgD-). Plasma cells were CD138+,plasmablasts were CD138+/B220-/CD19-.

For assessment of of either intracellular neuropeptides or neuropeptide receptors, cells were assessed from CD45+, live singlets and VIP or SP was stained intracellularly on CD3+, CD117+, Siglec F+, Ly6G+, or F4/80+ cells. VIP1R, MRGPRA1 was assessed on B220+/CD19+ or CD138+ cells.

For in vitro cultures cells were divided into B220+/CD19+ or CD138+. Then based on these gates assessed for IgG1 or IgE positivity. Ki67 was assessed from either CD138+/IgG1+ or CD138+/IgE+ gates. Exhaustion was assessed as CD11c+/CD23- from CD138+ and B220+/CD19+ gates.

### Immunofluorescence, histology, and microscopy

The lungs were perfused with PBS for immunostaining, followed by 4% paraformaldehyde (PFA) in PBS. Lungs were dissected and postfixed overnight in 4% PFA/PBS at 4 °C, incubated at 4 °C with 30% sucrose/PBS for two days, and stored in 0.1% sodium azide in PBS until cutting. Lungs were embedded in optimal cutting temperature compound (OCT, Tissue-Tek, PA), and 50-μm cryosections were cut at -20°C and then blocked for 4 h in PBS with 10% donkey serum, 2% bovine serum albumin (BSA) and 0.8% Triton X-100. Sections were immunostained with the following antibodies: B220 (Biolegend 103225; 1:100 FITC), CD45 (Invitrogen 14045182 1:200), E-cadherin (Thermoscientific MA512547 1:200), VIP (Thermoscientific PA578224, 1:200), Substance P (Thermoscientific PA5106934 1:200) with appropriate secondary antibodies (Goat anti mouse Cy3 Jackson Immuno 115-165-003; Goat anti rat Alexa 647 Jackson Immuno 112-605-003; Goat anti rabbit Alexa 488 Abcam AB150077), washed and mounted with prolong antifade diamond (Thermofisher P36961).

Vagal (nodose/jugular) and dorsal root ganglia were dissected, fixed in 4%PFA for 1 hour, cryopreserved in 30% sucrose overnight (all 4C), then placed in OCT, and 14µm sections were cut and placed on gelatin-coated slides. Sections were stained with TRPV1 (Alomone labs Clone- AB_2313819 1:200) followed by incubation with secondary antibody and secondary (Goat Anti Rabbit IgG H&L, Abcam). All sections were mounted with a Prolong diamond antifade compound (Thermofisher P36961).

For histology, lungs were embedded in paraffin and cut on a microtome (Leica) 5µm and stained with hematoxylin (Epredia 6765001) and eosin (Epredia 6766007), gram stain (Sigma HT90T- 1KT) or periodic acid, Schiff’s reagent (Sigma395B-1kit). Blood smears were stained with Giemsa solution (AB150670). Additionally, to confirm neutrophil counts in select experiments, Giemsa stain (Abcam AB150670) was used on lysed blood smears. Histological slides were mounted with permount toluene solution.

Sections were either imaged with a Leica SP8 confocal microscope or Leica Thunder system. Data were imaged and analyzed with Leica LASx software. For quantification of the proximity of B-cells to nerves, LASx online calipers were used to measure the distance from cell-to-nerve in 3D from the z-stack.

### B-cell culture

Spleens or lungs were used to isolate B-cells under sterile conditions. B-cells were isolated using the EasySep Mouse B-cell separation kit (StemCell Technologies 19854A). Cells were then treated in B-cell culture media made using 438 mL RPMI 1640 (Gibco 11835- 030), 2.5mL 1 M HEPES buffer (Sigma H3662), 5 mL GlutaMAX (Thermo Fisher Scientific 35050061), 5 mL penicillin/streptomycin stock (Sigma P0781), 30 mL heat-inactivated fetal bovine serum stock (Gibco A38400-01), and 454 μL β-mercaptoethanol (Gibco 21985-023) and vacuum filter sterilized (0.22 μm filter)^94^. Cells were then treated with either media + IL4 (20ng/ml, Sigma Aldrich I1020-5UG) + LPS (10µg/ml, L6529-1MG Sigma Aldrich), media+IL4+LPS+NPY (50ng/ml, Tocris 1153), media+IL4+LPS+VIP (50ng/ml, Tocris 1911) or, media+IL4+LPS+Substance P (50ng/ml, 1156 Tocris). After 96 hours media was stored at -80°C until immunoglobulin analysis. Cells were stained and analyzed with a Symphony A5 flow cytometer and data were processed with FlowJo (Treestar 10.881).

### Sample size and statistical analysis

We used animal numbers between 5 and 12 mice per experimental group/genotype for bacterial burden studies. For measurement of cytokine, neuropeptide, and immunoglobulins, 5-8 mice per group/genotype were used. For flow cytometry 5-10 mice per group/genotype were used. For in vitro experiments, at least 3 animals were pooled into one sample and 5-7 replicates were used. Bacterial burden data were analyzed with the two-way RM ANOVA (time x group) with either Tukey or Holm-Sidak posthoc test; FACS and cytokines were compared with ANOVA with Tukey or Holm-Sidak posthoc tests, two-tailed unpaired t-tests for parametric analyses, or Mann–Whitney test for nonparametric analyses. Correlations were run with Pearson correlation. Data were plotted in Prism (GraphPad).

## Acknowledgments

NJ is supported by NIH grant R21AI159221, R56AI175328, UCLA CTSI UL1TR001881-01 and T32KT4708 of the Regents of the University of California Tobacco-Related Diseases Research Program. T.A.D. is supported by NIH AI171795 and Veterans Affairs BLR&D BX005073.N.M. and D.A. are supported by a California Institute for Regenerative Medicine Stem Cell Biology Training Grant EDUC4-12837. The content is solely the responsibility of the authors and does not necessarily represent the official views of the National Institutes of Health. We are grateful to Dr. Isaac Chiu (Harvard) for providing TPV1-DTR mice as per the direction of Dr. Mark Hoon (NIH). We are grateful to Dr. Xin Sun for providing Vglut-Tdtomato mice. Figures 1a, 2g, 2h, 4a, 4p, 5a, 6a and Supplementary Figures 2a, 16a were made using Biorender.

## Author contributions

D.A. and F.Z. contributed to experimental design, conducted experiments and analyzed data. A.M. and N.M. contributed to experimental execution and data analysis. P.G. J.P., T. A.D, O.A. assisted with experimental design and analysis. M.S contributed to study design and manuscript preparation. N.J. designed the study, conducted experiments, analyzed data, prepared the manuscript and figures. All authors agree on the manuscript.

