## Supplementary Figures for "Sensory neurons regulate stimulus-dependent humoral immunity"

**Supplementary figure 1.**

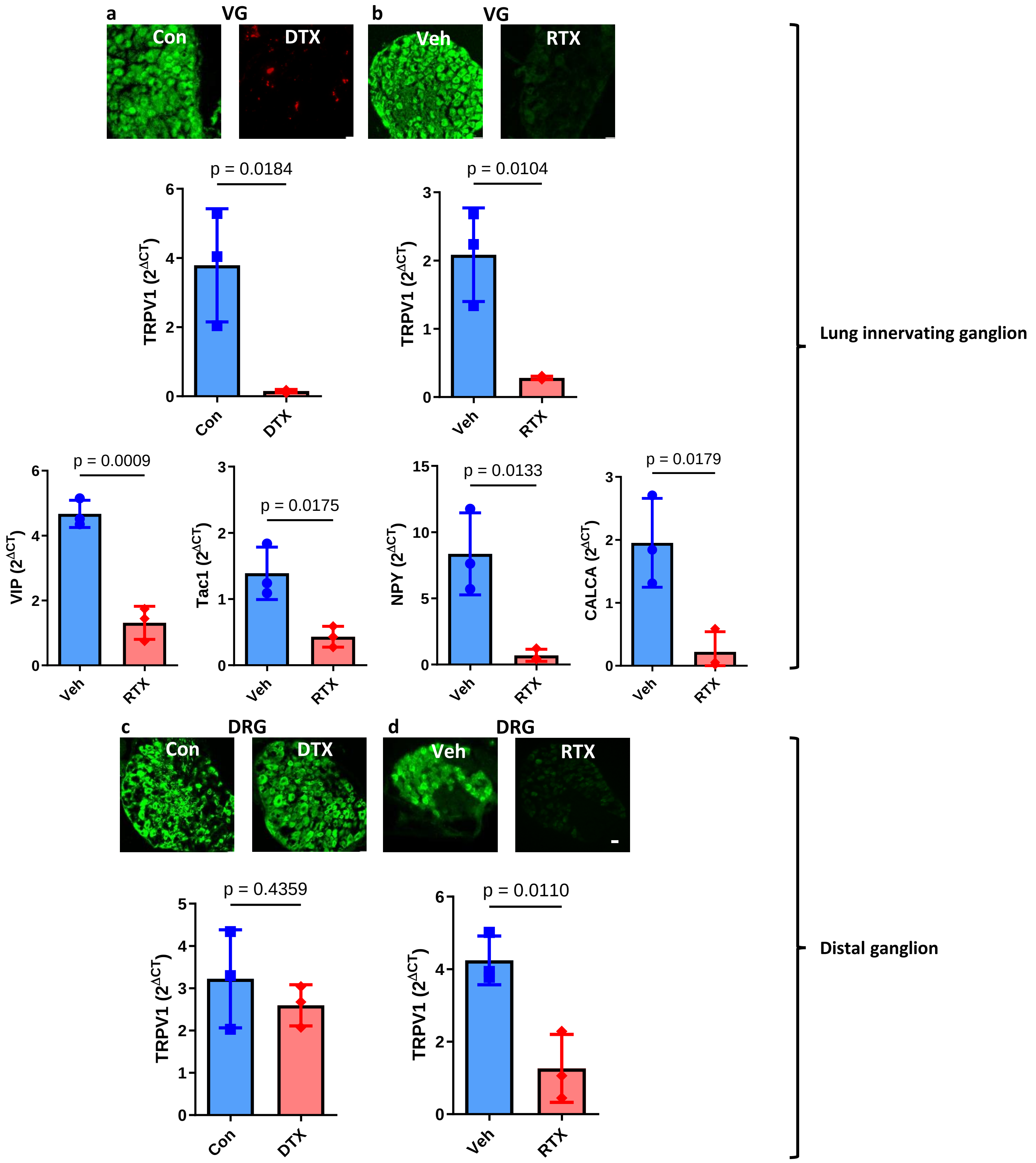

**Supplementary figure 1. TRPV1 knockdown by direct vagal (VG) diphtheria toxin (DTX) injection or global sensory neuron ablation with subcutaneous resiniferatoxin (RTX) injection. a)** qPCR demonstrates >85% downregulation of TRPV1 gene from vagal (VG) ganglia in DTX 200ng in 200nl VG injected versus sham injected TRPV1-DTR mice. TRPV1 is tagged with GFP in these mice, retrobeads were co-injected into DTX mice to demonstrate placement of injection as per Tränkner et al. In sham mice, the vagal ganglia were identified, and vehicle was injected in the vicinity of the VG. The prominent neuropeptides- vasoactive intestinal peptide (VIP), substance P (denoted by Tac1 gene), calcitonin gene related peptide (denoted by CALCA gene) and neuropeptide Y (NPY) were all down-regulated in the nodose/jugular ganglion with RTX treatment). **b)** TRPV1 gene is >85% downregulated in RTX compared to vehicle injected mice in VG. **c)** TRPV1 is maintained in dorsal root ganglia (DRG) of VG DTX injected mice compared to sham injected mice. **d)** TRPV1 is downregulated in DRG with global RTX injection. Two-sided t-test of ΔCT values. Housekeeping gene = hypoxyribosyltransferace. N=3 mice per sample (6 ganglia per sample), N=3 samples per group and condition.

**Supplementary figure 2.**

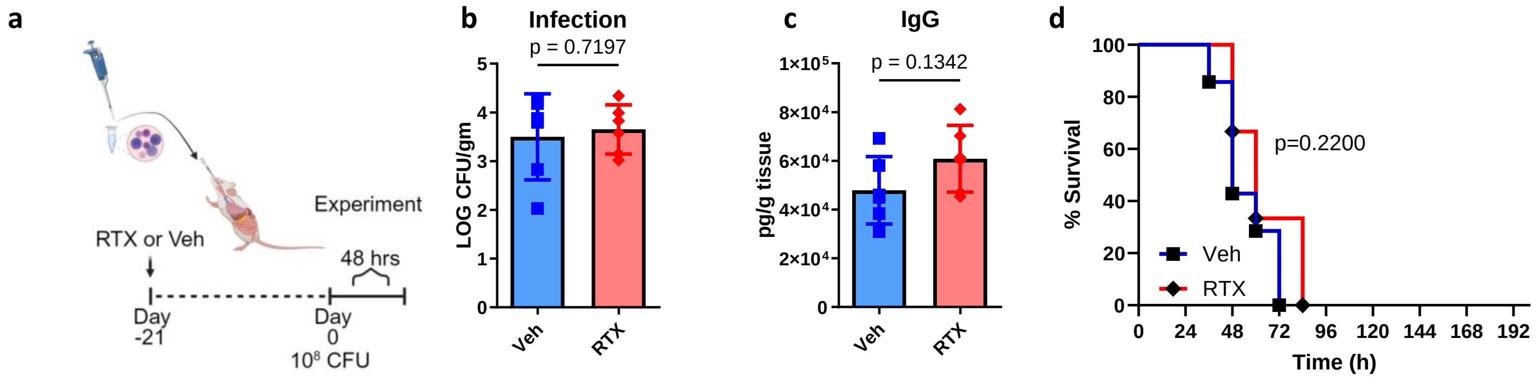

**Supplementary figure 2. Sensory neuron ablation does not alter bacterial clearance following acute infection with *S. pneumoniae*. a)** A single dose of $10^8$ CFU *S. pneumoniae (Serotype 19F, ATCC49619)* was delivered after treatment with vehicle or RTX as per our sensory neuron ablation model and then lungs were harvested to assess bacterial burden and immunoglobulin concentration 48h after infection. **b**) Log CFU per gram lung mass from acute infected sensory neuron intact (Veh) or sensory neuron ablated mice(RTX) show a similar bacterial burden. **c**) IgG concentrations from lung homogenates were similar between Veh and RTX 48h after acute infection. Veh: n=6, RTX: n=6. Two-sided T-test. **d**) A single dose of $10^6$ CFU *S. pneumoniae (Serotype 3, ATCC 6303) in 50µl* PBS was delivered after treatment with vehicle or RTX as per our sensory neuron ablation model and then survival was assessed. Veh: n=7, RTX: n=6. Data were compared with Log-rank (Mantel-Cox) test.

**Supplementary figure 3.**

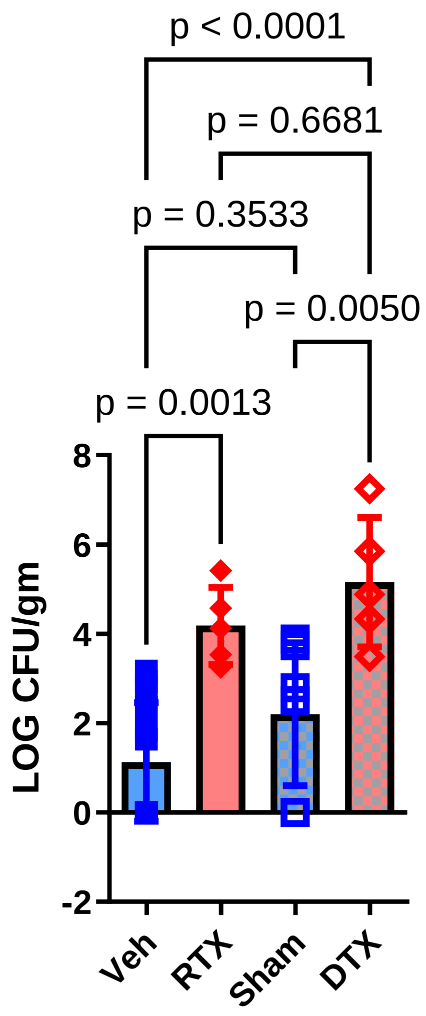

**Supplementary figure 3. The effect of bacterial clearance by sensory neurons is specific to vagal ganglia.** We compared the method of sensory neuron ablation on disease outcome. The bacterial burden in DTX or RTX mice was similarly increased 48h after the final infection with $10^8$ CFU of *S. pneumoniae (Serotype 19F, ATCC49619)* in comparison to Sham injection or Veh, respectively. Model of *S. pneumoniae* pre-exposure and infection as per Figure 1. Veh: n=13, RTX: n=5. Sham: n=7, DTX: n=5. One-way ANOVA with Tukey's post hoc test.

**Supplementary figure 4.**

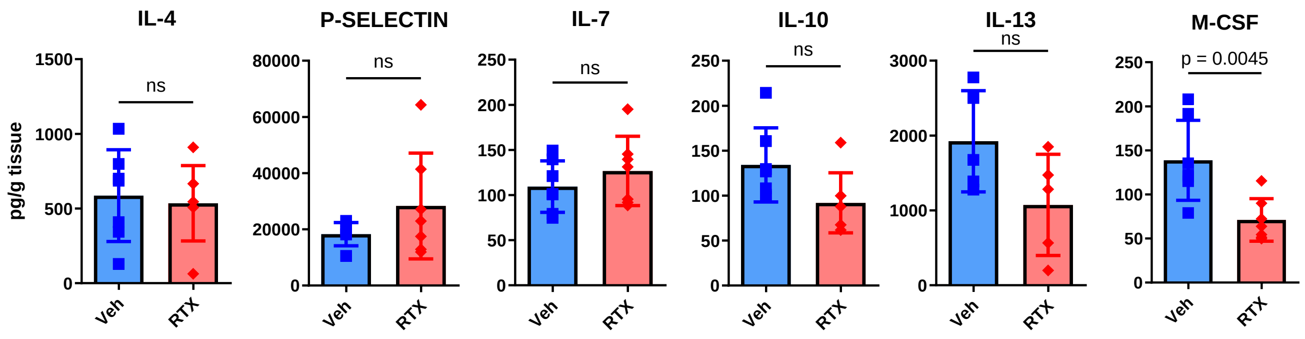

**Supplementary figure 4. Cytokines unaffected by sensory neuron ablation in response to pre-exposure and infection with *S. pneumoniae*.** Quantification of IL-4, P-Selectin, IL-7, IL-10, IL-13, M-CSF in sensory neuron intact (Veh, n=7) and sensory neuron ablated (RTX, n=6) mice 16h following the final $10^8$ CFU dose of *S. pneumoniae*. Two-sided t-test. Data were pooled from 2 independent experiments.

Supplementary figure 5.

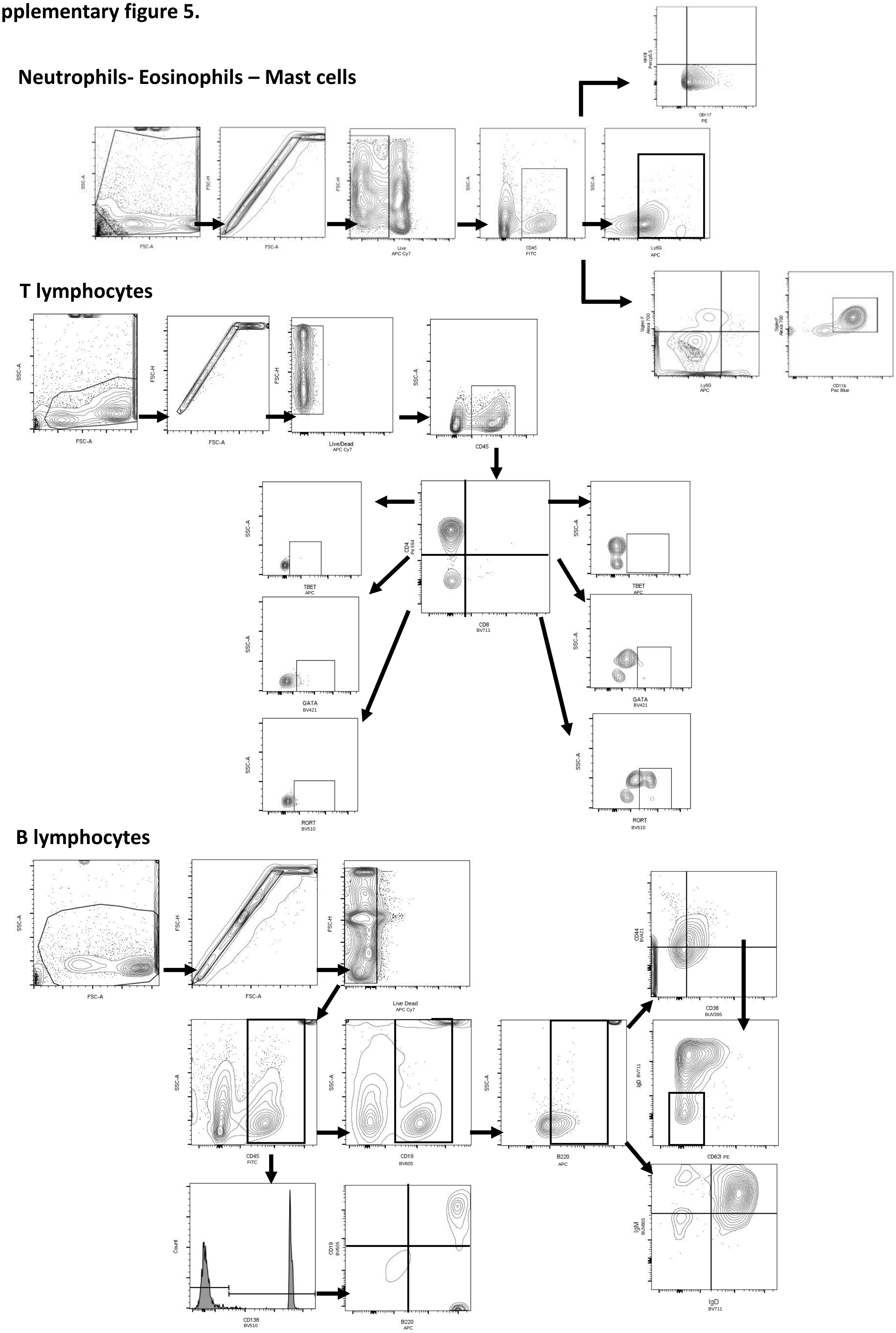

**Supplementary Figure 5. Gating strategy for Neutrophils, Eosinophils, Mast cells, T lymphocytes and B-lymphocytes.** Gating used for in vivo flow cytometry data from lungs, spleen and bone marrow collected on BD Symphony A5.

**Supplementary figure 6.**

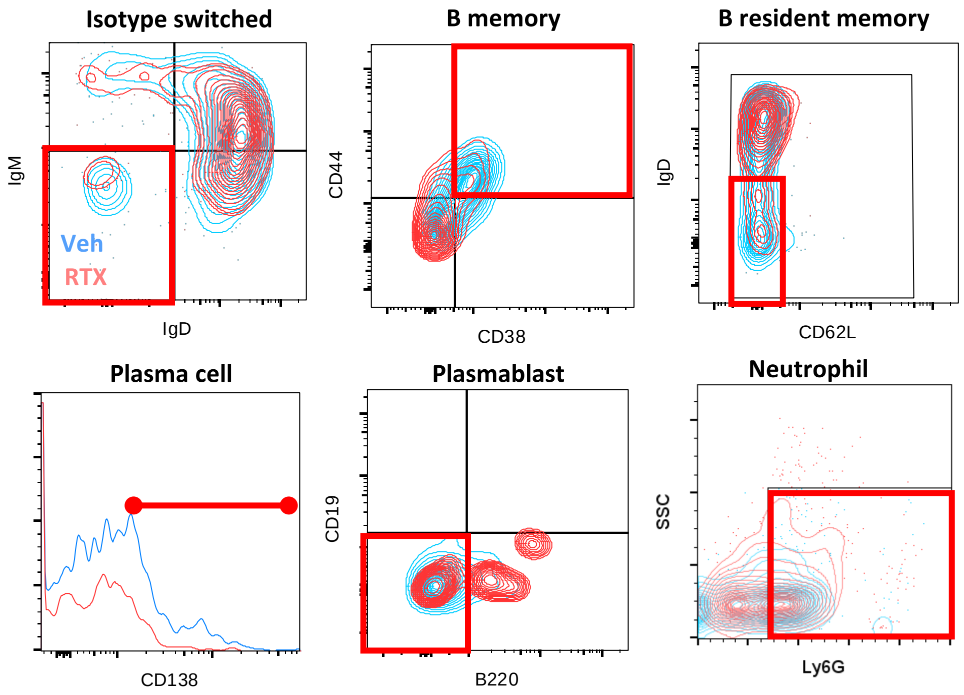

**Supplementary Figure 6. Comparisons B-cell populations between sensory neuron intact and depleted mice at 48h after final infection which was preceded with pre-exposure.** Gating used for *in vivo* flow cytometry data for B-cell populations. Vehicle treated sensory neuron intact mice (Veh) are in blue and sensory neuron depleted mice (RTX) are in red. The representative gate for Isotype Switched, B memory, B resident memory, Plasma cell, Plasmablast and Neutrophils are shown in red gates. Populations were gated as per Supplementary Figure 5. Collected on BD Symphony A5.

**Supplementary figure 7.**

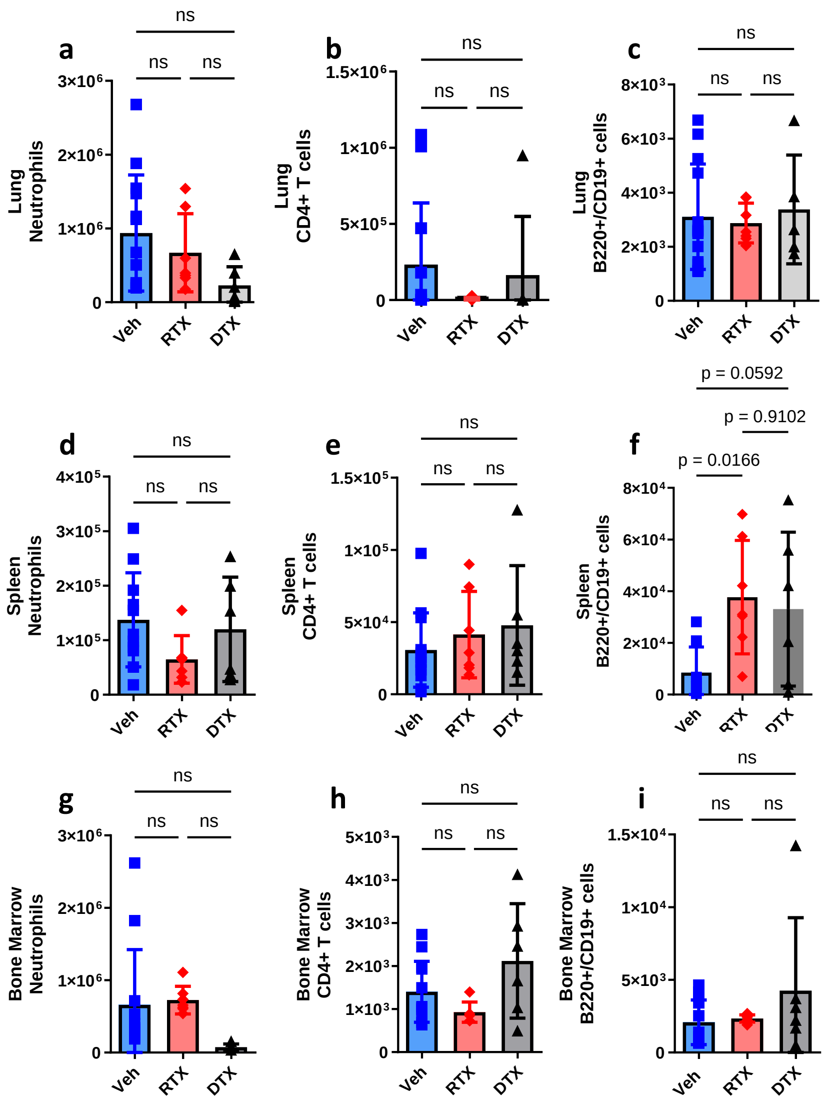

**Supplementary Figure 7. Sensory neuron ablation does not effect bone marrow or spleen cells. a-c)** Lung, **d-f)** Spleen and **g-i)** Bone Marrow populations of **a, d, f)** Neutrophils (CD45+/Ly6G+), **b, e, h)** T helper lymphocytes (CD45+/CD4+/CD8-) or **c, f, i)** B-lymphocytes (CD45+/B220+/CD19+ all gated on live/singlets) were maintained in both conditions 16h following pre-exposure and infection with *Streptococcus Pneumoniae 19F*. DTX and RTX models showed similarities with cell populations. Total cell populations were determined with counting beads. Veh: n=13, RTX: n=7, DTX: n=6. Data from 3 independent experiments. One-way ANOVA with Tukey's post hoc test.

**Supplementary figure 8.**

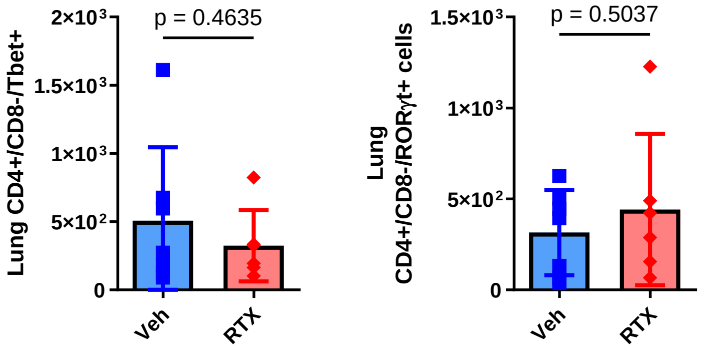

**Supplementary figure 8. Sensory neuron ablation does not reduce T-lymphocytes 48h after infection with *S. pneumoniae*.** T helper cells were similar in sensory neuron intact and ablated mice. Vehicle: n=6, RTX: n=6. Cells were gated on live/CD45+ cells. Counting beads determined cell populations. Data from 2 independent experiments. Two-sided t-test.

**Supplementary figure 9.**

a

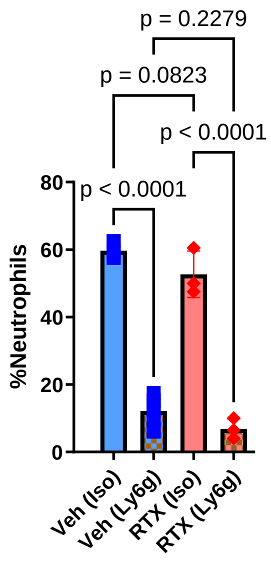

b

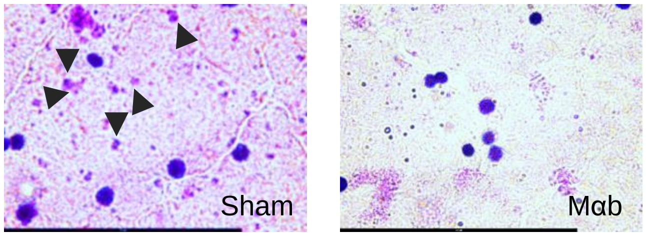

**Supplementary figure 9. Neutrophil depletion. a**) Circulating neutrophils were suppressed with anti-1A8 antibody. Veh (Iso): n=7, RTX (Iso):n=3, Veh (Ly6G): n=8, RTX (Ly6G): n=3. Wight-Giemsa stain on blood smear. One-way ANNOVA with Tukey's post-hoc test. Data from 2 independent experiments. **b**) Representative blood smears from Isotype treated and neutrophil depletion treated mice. Arrowheads point to neutrophils. Scalebar=50um.

**Supplementary figure 10.**

**a**

| | CFU | NPY | SubP | CGRP | VIP | IgA | IgM | IgE | IgG1 | IgG2a | IgG2b | IgG3 |
|---|---|---|---|---|---|---|---|---|---|---|---|---|
| **CFU** | 1.00 | -0.67 | -0.58 | -0.36 | -0.76 | -0.60 | -0.65 | -0.56 | -0.85 | -0.52 | -0.73 | -0.70 |
| **NPY** | -0.67 | 1.00 | 0.77 | 0.63 | 0.77 | 0.73 | 0.52 | 0.60 | 0.80 | 0.59 | 0.80 | 0.88 |
| **SubP** | -0.58 | 0.77 | 1.00 | 0.81 | 0.81 | 0.87 | 0.55 | 0.36 | 0.67 | 0.30 | 0.70 | 0.54 |
| **CGRP** | -0.36 | 0.63 | 0.81 | 1.00 | 0.55 | 0.74 | 0.53 | 0.01 | 0.64 | 0.22 | 0.39 | 0.47 |
| **VIP** | -0.76 | 0.77 | 0.81 | 0.55 | 1.00 | 0.86 | 0.62 | 0.32 | 0.72 | 0.34 | 0.61 | 0.59 |
| **IgA** | -0.60 | 0.73 | 0.87 | 0.74 | 0.86 | 1.00 | 0.63 | 0.17 | 0.77 | 0.28 | 0.65 | 0.51 |
| **IgM** | -0.65 | 0.52 | 0.55 | 0.53 | 0.62 | 0.63 | 1.00 | 0.08 | 0.74 | 0.64 | 0.38 | 0.56 |
| **IgE** | -0.56 | 0.60 | 0.36 | 0.01 | 0.32 | 0.17 | 0.08 | 1.00 | 0.41 | 0.57 | 0.83 | 0.63 |
| **IgG1** | -0.85 | 0.80 | 0.67 | 0.64 | 0.72 | 0.77 | 0.74 | 0.41 | 1.00 | 0.62 | 0.74 | 0.84 |
| **IgG2a** | -0.52 | 0.59 | 0.30 | 0.22 | 0.34 | 0.28 | 0.64 | 0.57 | 0.62 | 1.00 | 0.56 | 0.80 |
| **IgG2b** | -0.73 | 0.80 | 0.70 | 0.39 | 0.61 | 0.65 | 0.38 | 0.83 | 0.74 | 0.56 | 1.00 | 0.73 |
| **IgG3** | -0.70 | 0.88 | 0.54 | 0.47 | 0.59 | 0.51 | 0.56 | 0.63 | 0.84 | 0.80 | 0.73 | 1.00 |

**b**

| | CFU | NPY | SubP | CGRP | VIP | IgA | IgM | IgE | IgG1 | IgG2a | IgG2b | IgG3 |
|---|---|---|---|---|---|---|---|---|---|---|---|---|
| CFU | | 0.01653 | 0.04710 | 0.24975 | 0.00438 | 0.03758 | 0.02219 | 0.06014 | 0.00045 | 0.08170 | 0.00667 | 0.01060 |
| NPY | 0.01653 | | 0.00326 | 0.02953 | 0.00320 | 0.00762 | 0.08230 | 0.04060 | 0.00180 | 0.04487 | 0.00194 | 0.00013 |
| SubP | 0.04710 | 0.00326 | | 0.00130 | 0.00142 | 0.00022 | 0.06640 | 0.25111 | 0.01760 | 0.33736 | 0.01195 | 0.06704 |
| CGRP | 0.24975 | 0.02953 | 0.00130 | | 0.06616 | 0.00577 | 0.07582 | 0.97280 | 0.02450 | 0.50163 | 0.21350 | 0.12286 |
| VIP | 0.00438 | 0.00320 | 0.00142 | 0.06616 | | 0.00030 | 0.03329 | 0.31057 | 0.00884 | 0.27689 | 0.03333 | 0.04195 |
| IgA | 0.03758 | 0.00762 | 0.00022 | 0.00577 | 0.00030 | | 0.02699 | 0.60501 | 0.00341 | 0.38203 | 0.02330 | 0.09246 |
| IgM | 0.02219 | 0.08230 | 0.06640 | 0.07582 | 0.03329 | 0.02699 | | 0.80710 | 0.00605 | 0.02400 | 0.22491 | 0.05849 |
| IgE | 0.06014 | 0.04060 | 0.25111 | 0.97280 | 0.31057 | 0.60501 | 0.80710 | | 0.18249 | 0.05487 | 0.00080 | 0.02906 |
| IgG1 | 0.00045 | 0.00180 | 0.01760 | 0.02450 | 0.00884 | 0.00341 | 0.00605 | 0.18249 | | 0.03256 | 0.00545 | 0.00066 |
| IgG2a | 0.08170 | 0.04487 | 0.33736 | 0.50163 | 0.27689 | 0.38203 | 0.02400 | 0.05487 | 0.03256 | | 0.05679 | 0.00159 |
| IgG2b | 0.00667 | 0.00194 | 0.01195 | 0.21350 | 0.03333 | 0.02330 | 0.22491 | 0.00080 | 0.00545 | 0.05679 | | 0.00709 |
| IgG3 | 0.01060 | 0.00013 | 0.06704 | 0.12286 | 0.04195 | 0.09246 | 0.05849 | 0.02906 | 0.00066 | 0.00159 | 0.00709 | |

**Supplementary figure 10. Neuropeptides and immunoglobulins correlate with bacterial burden.** Lung bacterial burden (Figure 1b- 48h timepoint), immunoglobulins (Figure 2a) and neuropeptides (Figure 3b, c, d, e) were compared with Pearson correlation. CFU was inversely correlated with neuropeptides and immunoglobulins with VIP and IgG1 showing the strongest relationships to Lung bacterial burden (CFU). Neuropeptides and immunoglobulins were positively correlated with one another. **a**) Pearson r. **b**) P-values.

**Supplementary figure 11.**

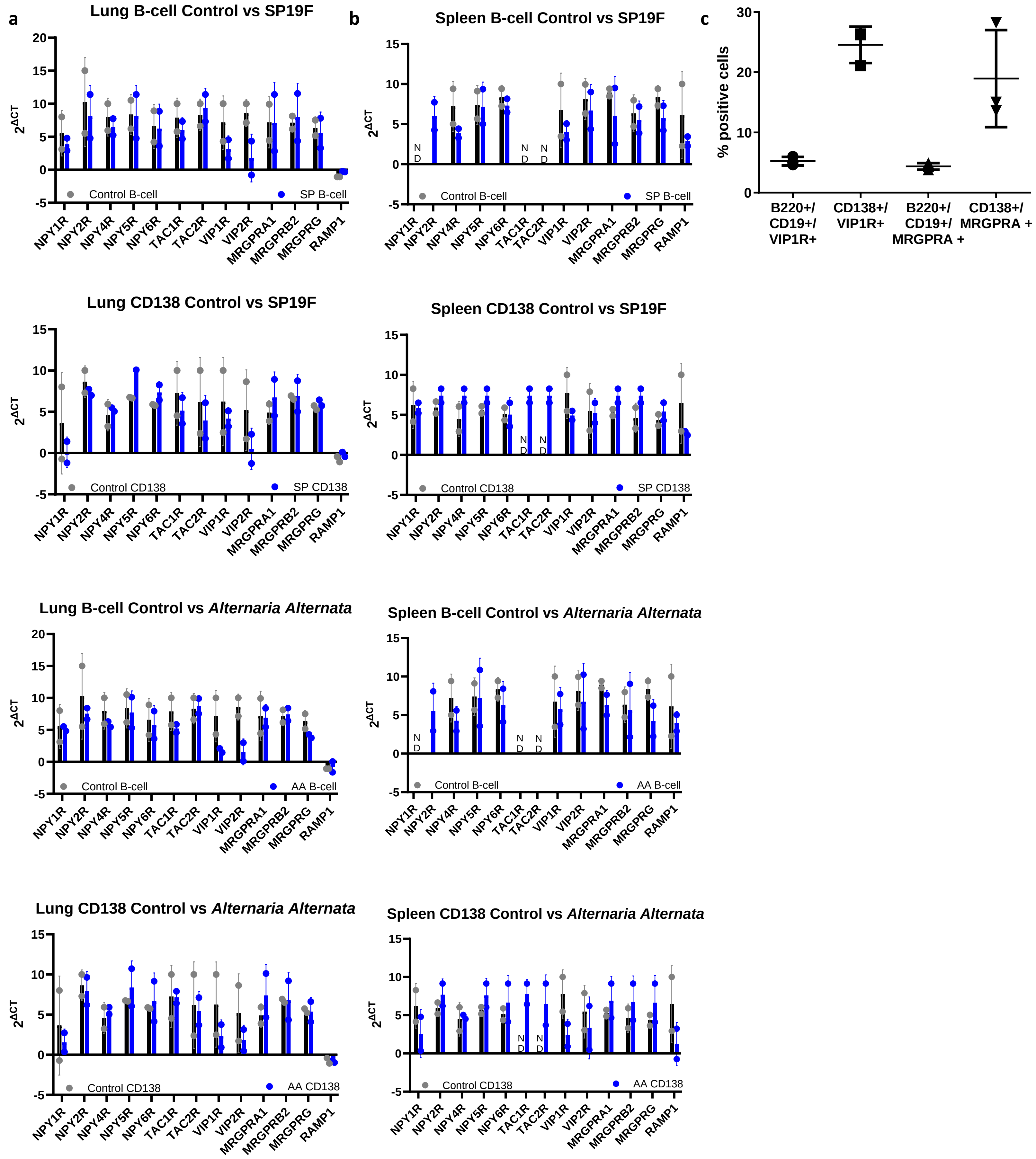

**Supplementary figure 11. Prominent neuropeptide receptors in isolated B-cells.** CD138+ and B-cells were isolated from the lungs (**a**) and spleen (**b**) of naïve, S. pneumoniae and A. Alternata mice (n=5 mice were pooled per sample, 2 samples per group). Cells were isolated with the CD138 positive or B-cell negative EasySep selection kit. qPCR for prominent neuropeptide receptors was run, and data were expressed as ΔCT from the hypoxyribosyltransferase housekeeping gene. No differences between groups within the respective subset were demonstrated for any receptor. Although we note that VIP1R, VIP2R, RAMP1 are highly expressed in all samples. **c**) Protein surface expression from splenic naïve isolated B- cells were gated on Live B220+/CD19+ or CD138+ cells. VIP1R and MRGPRG were highly expressed on plasma cells. N=3.

**Supplementary figure 12.**

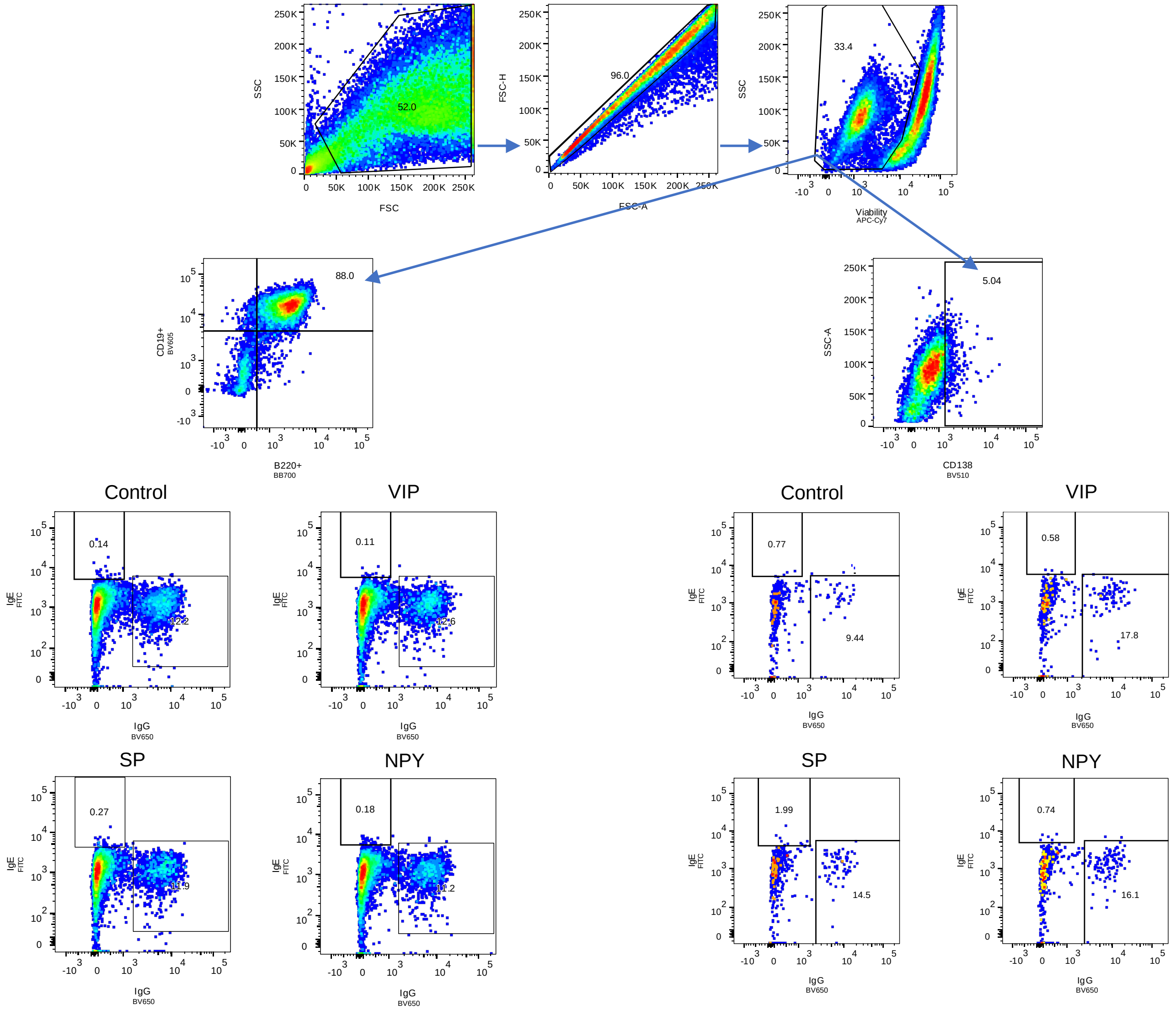

**Supplementary Figure 12. Comparisons of neuropeptide stimulated cultured B-cells.** Gating used for *in vitro* flow cytometry data for B-cell populations. Collected on BD Symphony A5.

**Supplementary figure 13.**

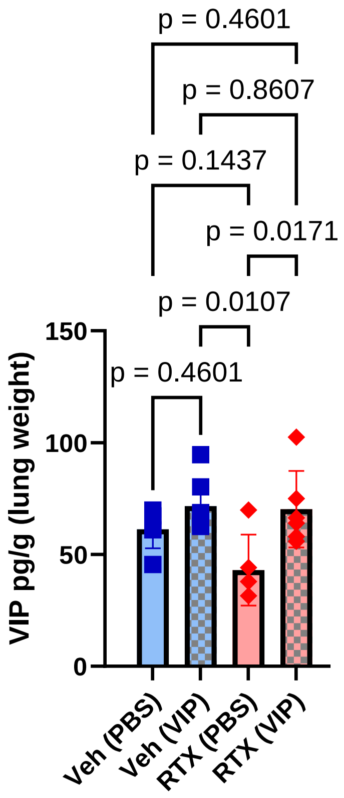

**Supplementary figure 13. Vasoactive intestinal peptide supplementation increases vasoactive intestinal peptide in RTX mice.** Mice were treated as per Figure 4a. VIP was measured from lung homogenates and expressed per lung weight. VIP supplementation significantly increased VIP levels in lungs of RTX mice 16hrs after final infectious dose of bacteria. Veh (PBS): n=6, Veh (VIP): n=7, RTX(PBS): n=5, RTX(VIP): n=6. One-way ANOVA with Holm sidak post hoc test. Data from 3 independent experiments.

**Supplementary figure 14.**

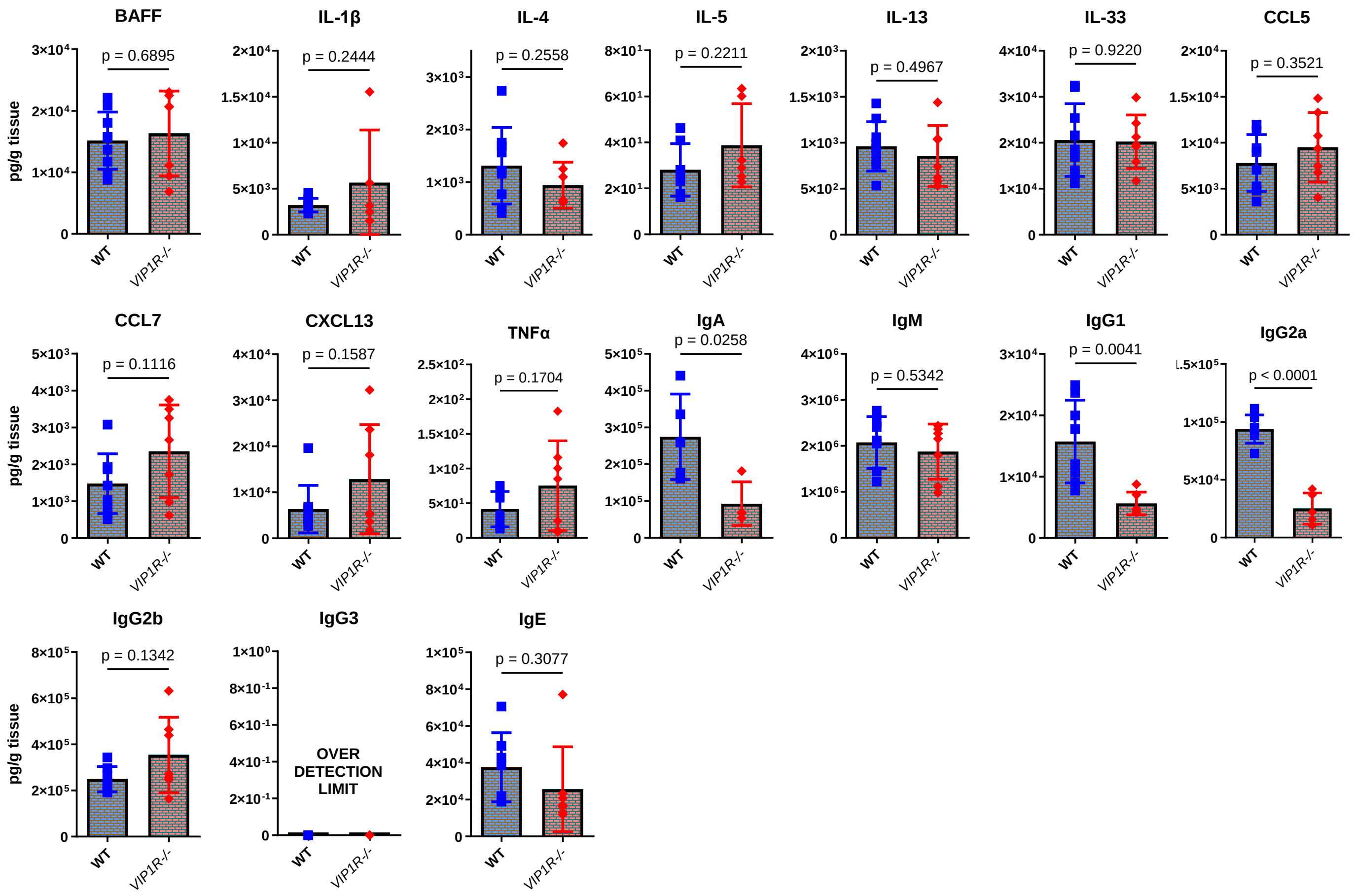

**Supplementary figure 14. VIP 1 receptor knockout mice have reduced immunoglobulins, but not cytokines compared to WT mice following pre-exposure and infection to *S. pneumoniae*.** Quantification of BAFF, IL-1β, IL-4, IL-5, IL-13, IL-33, CCL5, CCL7, CXCL13, TNFα, IgA, IgM, IgG1, IgG2a, IgG2b, IgG3, IgE in WT and VIP 1 receptor knockout mice (VIP1R-/-) following pre-exposure and infection to *S. pneumoniae* WT: n=6-8, VIP1R-/-: n=5-7. Two-sided t-test. Data from 2 independent experiments.

**Supplementary figure 15.**

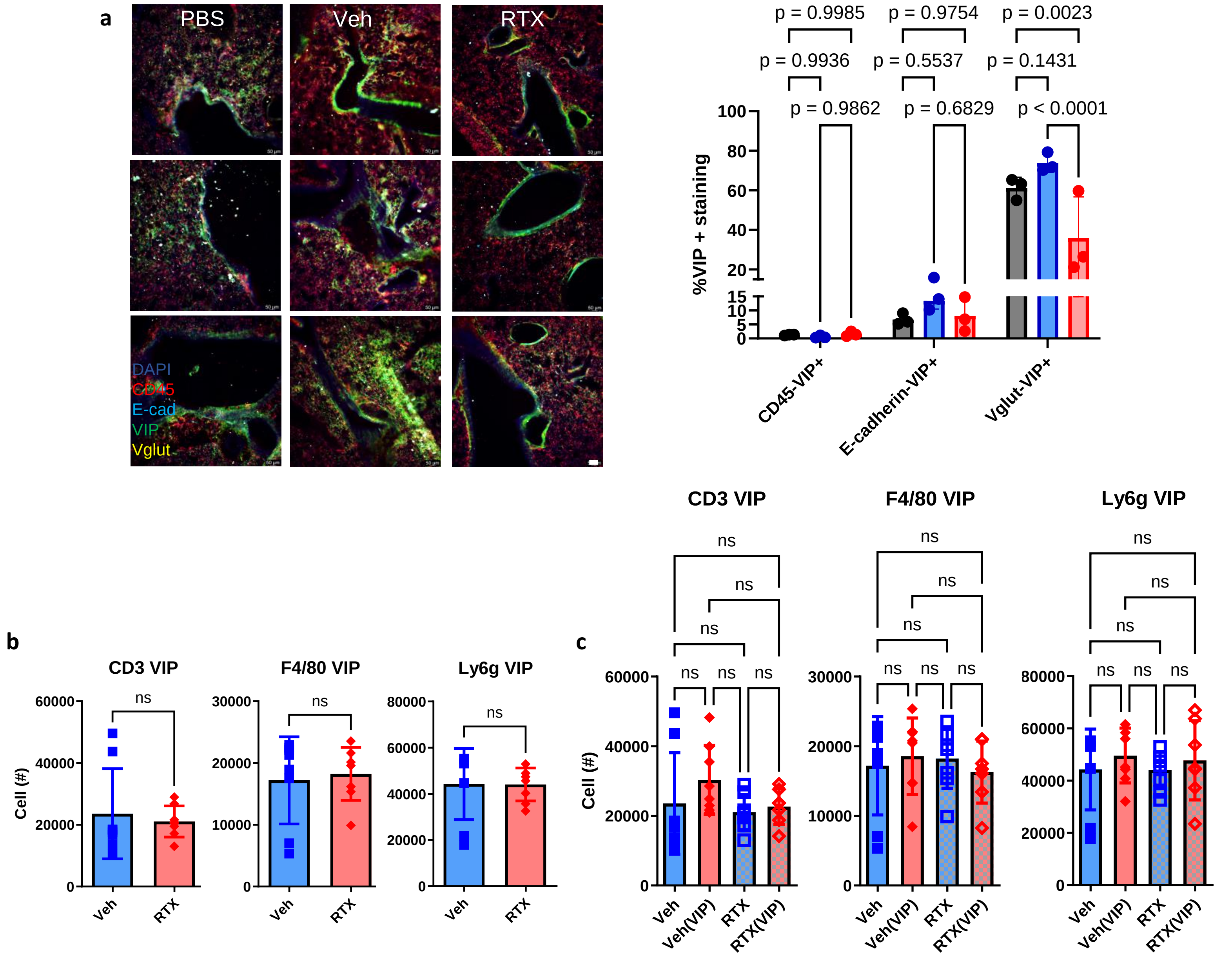

**Supplementary figure 15. VIP from non-neuronal sources is similar between vehicle and RTX infected mice. a**) Vglutcre-tdtomato mice underwent our pre-exposure/infection model as per Figure 1a. Lungs were harvested and stained for CD45 (leukocytes), E-cadherin (epithelium) and VIP. The VIP fluorescence signal was normalized to the CD45, E-cadherin or Vglut fluorescence signal. RTX only reduced VIP in Vglut expressing cells. Two-way ANOVA with Holm-Sidak post-hoc test. N=3 per group. Scalebar= 50µm. **b**) Lungs from Veh or RTX mice were processed for flow cytometry. Cells were gated from CD45 live, singlets. RTX and Vehicle exposed to our pre-exposure and infection model as per Figure 1 had similar VIP expression in T-cells (CD3), Macrophages (F4/80) and neutrophils (Ly6g). N=8 per group. Two-sided t-test. **c**) Veh and RTX mice underwent VIP supplementation as per Figure 4a. Cells were stained as per panel b. No differences were demonstrated between groups for VIP from leukocytes. N=7-8 per group, One-way ANOVA with Holm-Sidak post-hoc test. Two independent experiments.

**Supplementary figure 16.**

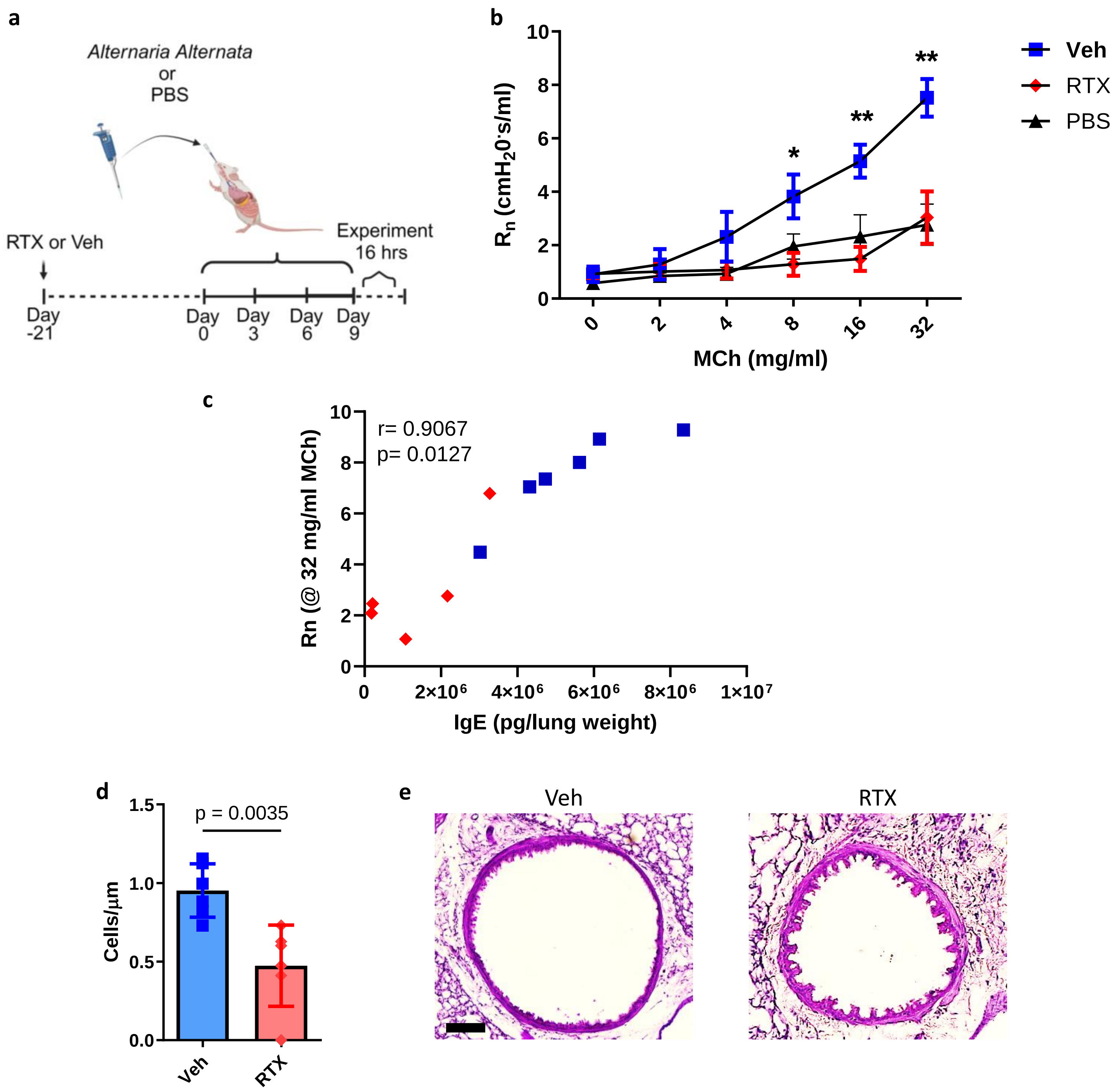

**Supplementary figure 16. Sensory neuron ablation reduces airway hyperresponsiveness and goblet cell metaplasia in *Alternaria Alternata* induced asthma. a)** *A. alternata* induced asthma was elicited as per Cavagnero et al. 25µg of *A. alternata* extract in 50µl PBS (PBS only for veh) was delivered as per the schematic. Experiments took place 16hrs later. **b**) RTX treated mice had a response to doubling doses of methacholine similar to PBS control mice and reduced compared to sensory neuron intact mice treated with *A. alternata*. PBS: n=5, Veh: n=6, RTX: n=5. Two-way ANOVA with Tukey's post hoc test. * indicates difference between Veh and RTX. ** indicates difference from Veh and RTX, Veh and PBS. Data from 2 independent experiments. **c)** Correlation of IgE (from Figure 5b) and Rn @ MCh (32mg/ml) **d)** Goblet cell metaplasia (Cells/µm) are reduced in RTX compared to sensory neuron intact *A alternata* treated mice. Veh: n=6, RTX: n=7.Two-sided t-test. Data from 2 independent experiments. **e**) Representative images of periodic acid Schiff's reagent stain to visualize mucous producing Goblet cells. Scalebar=20um.

**Supplementary figure 17.**

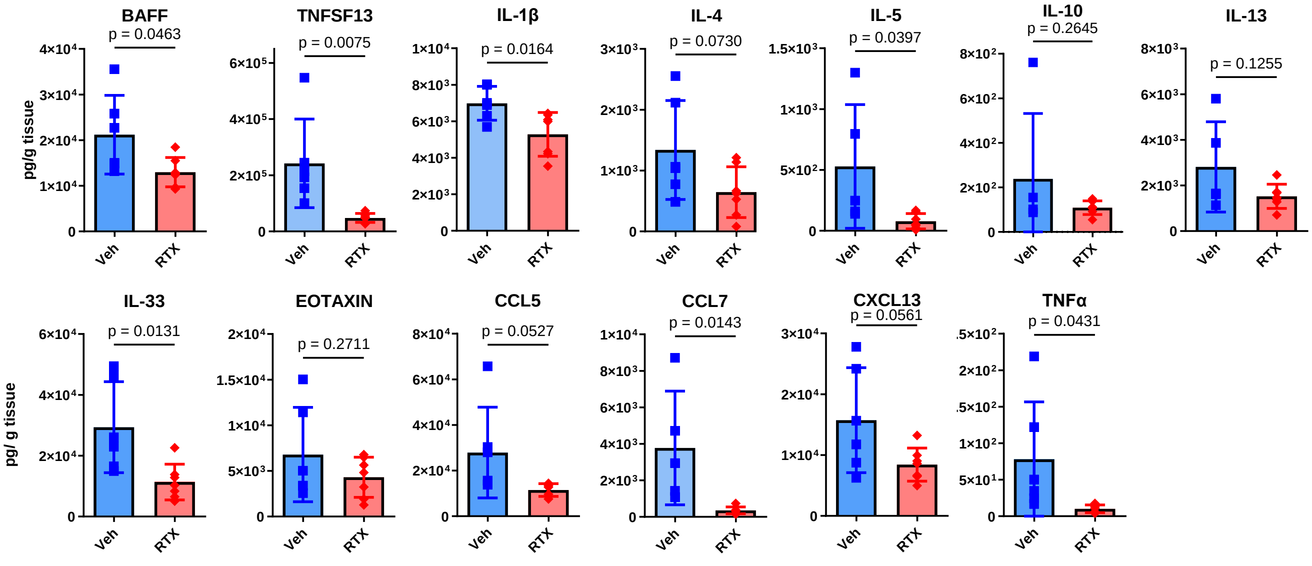

**Supplementary figure 17. Sensory neuron ablation suppresses inflammatory, b-cell survival and recruiting cytokines and chemokines and Th2 cytokines following *A. alternata treatment*.** Quantification of BAFF, TNFSF13, IL-1β, IL-4, IL-5, IL-10, IL-13, IL-33, Eotaxin, CCL5, CCL7, CXCL5 and, TNFα in sensory neuron intact and sensory neuron ablated (RTX) after the final dose of *A. alternata*. Veh: n=5-7, RTX: n=6-7. Two-sided t-test. Data from 3 independent experiments.

**Supplementary figure 18.**

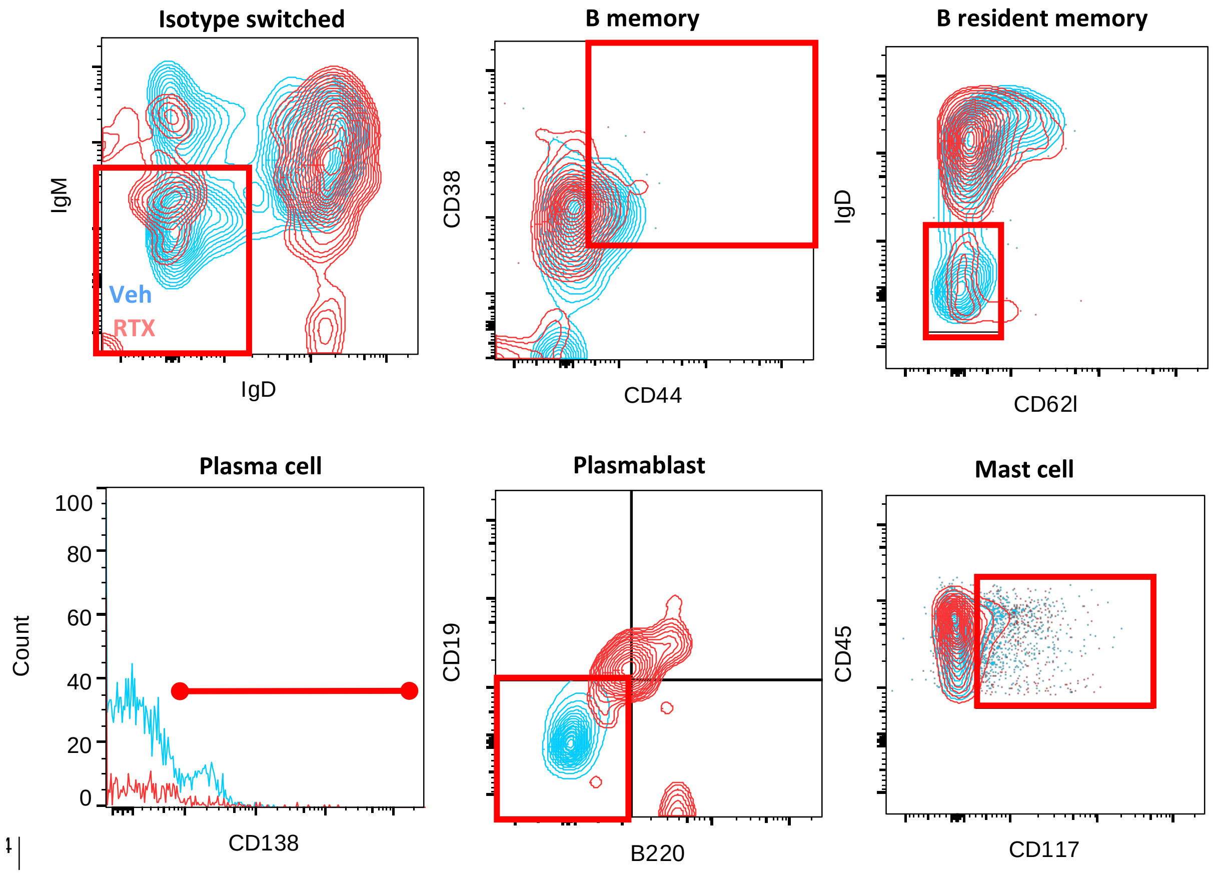

**Supplementary Figure 18. Comparisons between sensory neuron intact and depleted mice with *Alternaria Alternata* asthma 16h after the last dose of *Alternaria Alternata*.** Gating used for in vivo flow cytometry data for B-cell populations. Vehicle treated sensory neuron intact mice are in blue and RTX sensory neuron depleted mice are in red. The representative gate for Isotype Switched, B memory, B resident memory, Plasma cell, Plasmablast and Mast cells are shown and gated as per Supplementary Figure 5. Collected on BD Symphony A5.

**Supplementary figure 19.**

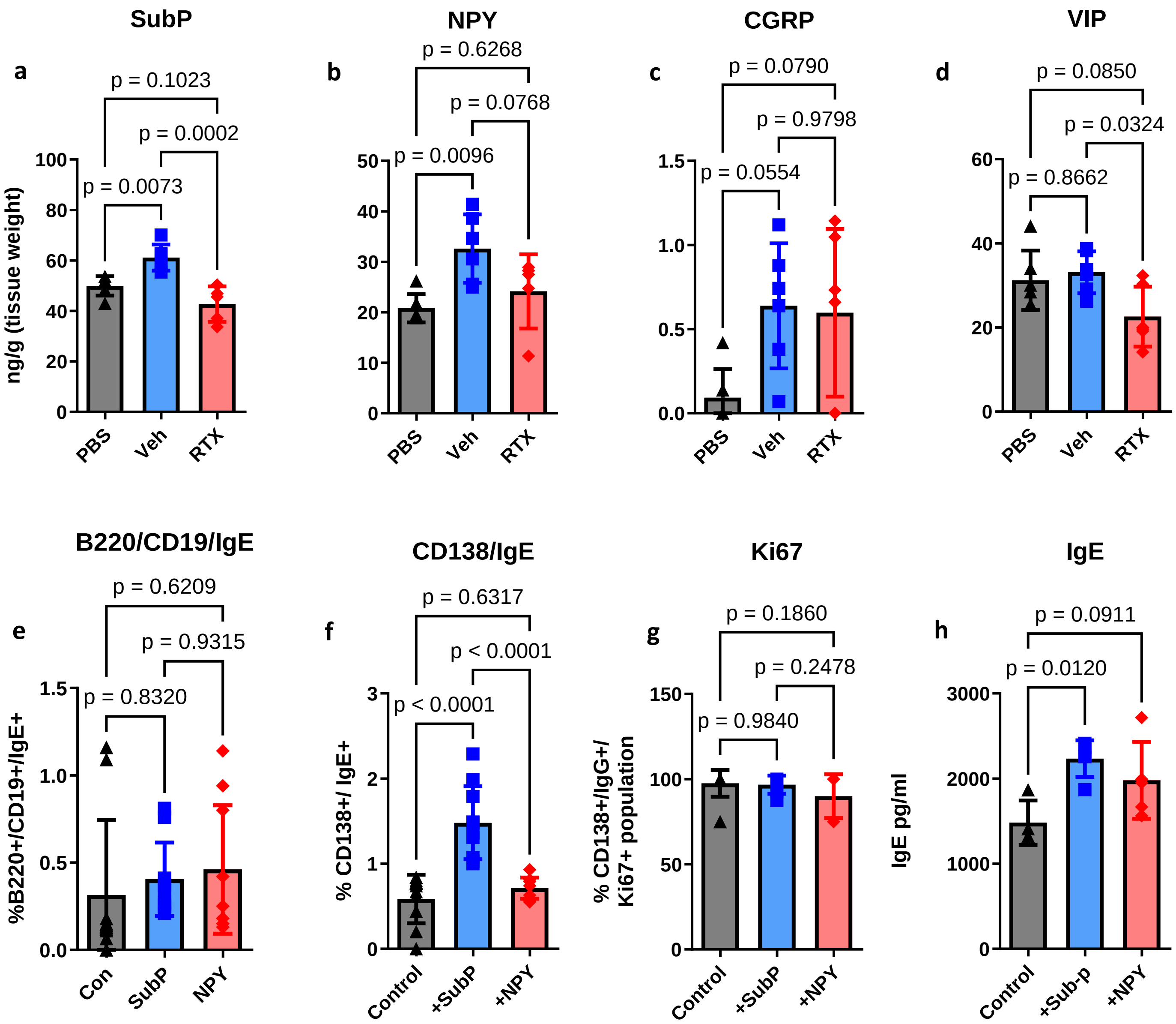

**Supplementary figure 19. *A. alternata* increases Substance P which augments IgE production.** *A. alternata* increased lung concentrations of Substance P (**a**) and NPY (**b**). CGRP (**c**) and VIP (**d**) were not significantly increased following *A. alternata* induction. Substance P (**a**) and VIP (**d**) were significantly reduced with RTX treatment. Analyzed from lung homogenates with ELISA. PBS: n=6, Veh: n=6, RTX: n=6. One way ANOVA with Tukey's post-hoc test. Data from 3 independent experiments. **e-h)** B cells were isolated from spleens of naïve mice and cultured in media with IL4+LPS, and supplemented with Substance P (SubP) or NPY. Cells and media were sampled after 96 hours of incubation. **e**) Substance P did not increase bound IgE on B220/CD19+ cells. **f**) Substance P significantly increased bound IgE in CD138+ plasma cells. **g)** Proliferation as assessed by intracellular Ki67 staining was not affected by neuropeptides (on CD138+ cells). **h**) Secreted IgE was increased by Substance P preferentially. IL4+LPS: n=4, +SubP: n=5, +NPY: n=5. One-way ANOVA and Tukey's post-hoc test. Data from 2 independent experiments.

**Supplementary figure 20.**

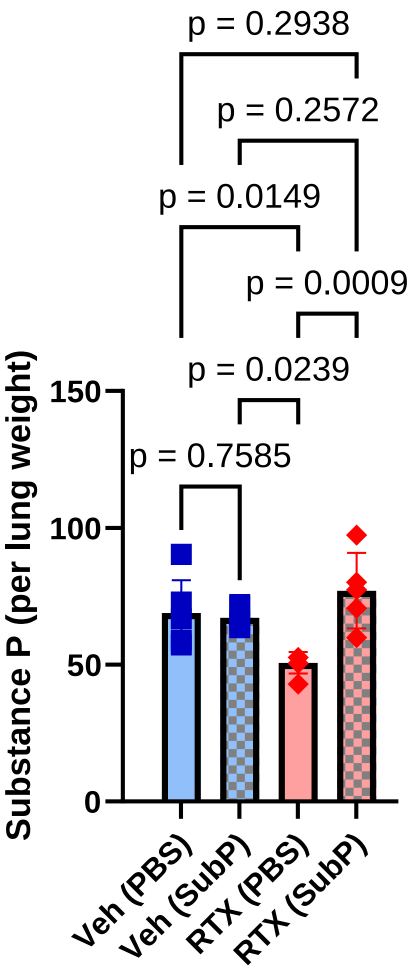

**Supplementary figure 20. Substance P supplementation increases Substance P in RTX mice.** Mice were treated as per Figure 6a. Substance P was measured from lung homogenates and expressed per lung weight. Substance P supplementation significantly increased Substance P levels in lungs of RTX mice 16hrs after final *A. alternata* dose. Veh (PBS): n=6, Veh (SubP): n=6, RTX(PBS): n=6, RTX(SubP): n=5. One-way ANOVA with Holm sidak post hoc test. Data from 3 independent experiments.

**Supplementary figure 21.**

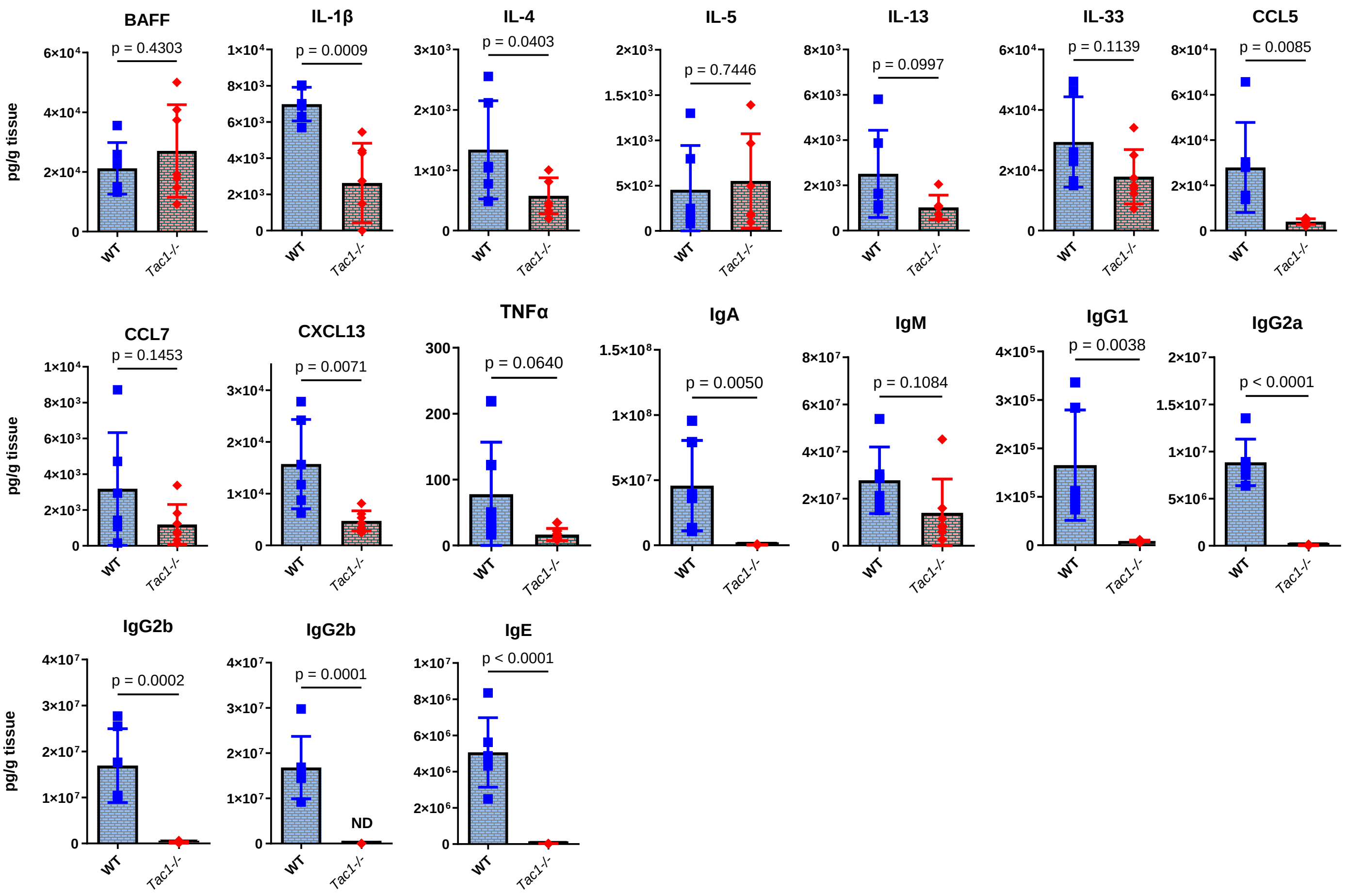

**Supplementary figure 21. Tac1 knockout mice have reduced inflammatory, b-cell survival, recruiting cytokines, chemokines, Th2 cytokines and immunoglobulins following *A. alternata treatment*.** Quantification of BAFF, IL-1β, IL-4, IL-5, IL-13, IL-33, CCL5, CCL7, CXCL13, TNFα, IgA, IgM, IgG1, IgG2a, IgG2b, IgG3, IgE (repeated from figure 6) in Wild type and Tac1 (substance P gene) -/- mice after the final dose of *A alternata*. WT: n=7-8, Tac1KO: n=7-8. Two-sided t-test. Data from 2 independent experiments.

**Supplementary figure 22.**

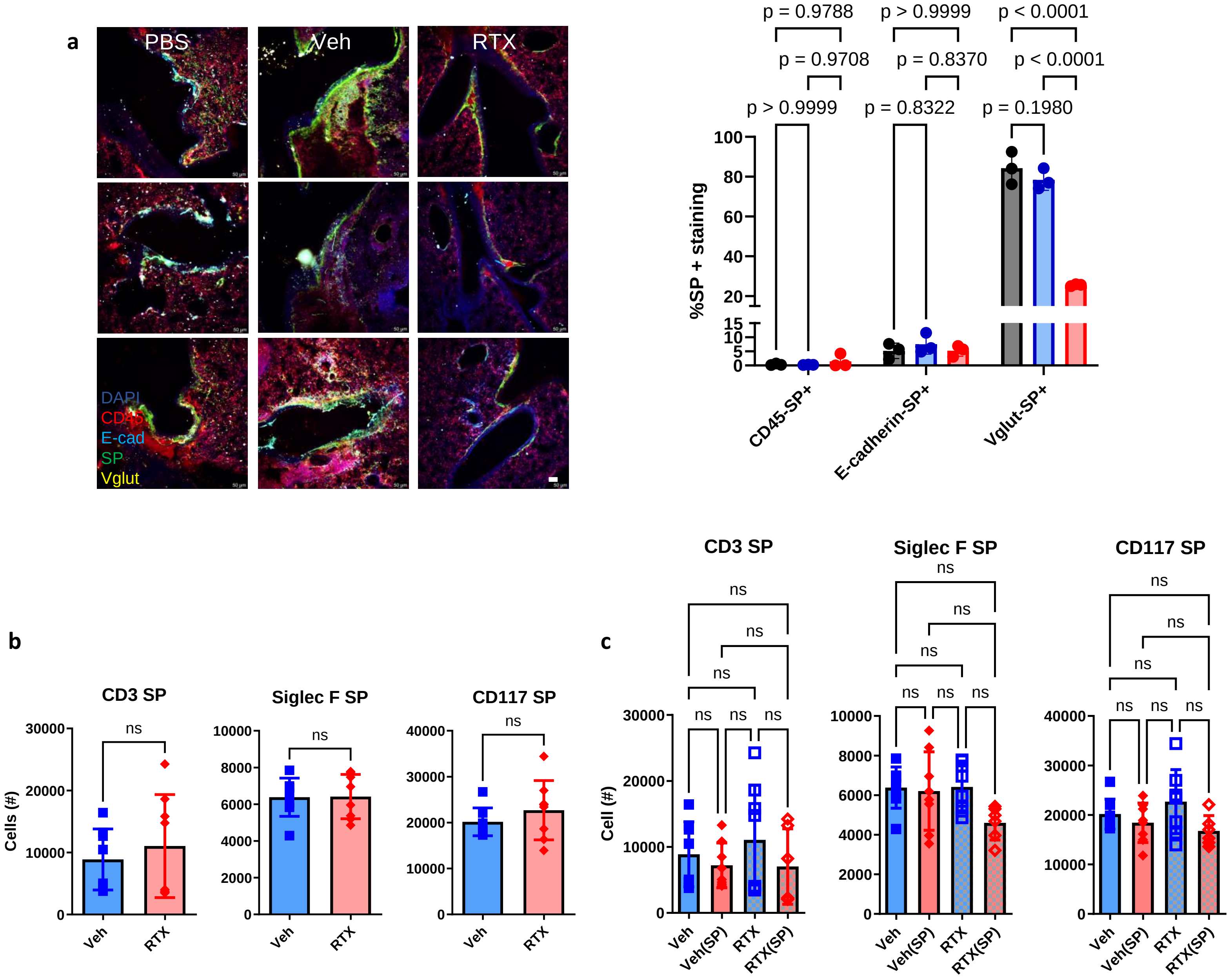

**Supplementary figure 22. Substance P from non-neuronal sources is similar between vehicle and RTX *A. alternata* mice. a**) Vglutcre-tdtomato mice underwent A Alternata as per Figure 5. Lungs were harvested and stained for CD45 (leukocytes), E-cadherin (epithelium) and Substance P (SP). The SP fluorescence signal was normalized to the CD45, E-cadherin or Vglut fluorescence signal. RTX only reduced VIP in Vglut expressing cells. Two-way ANOVA with Holm-Sidak post-hoc test. N=3 per group. Scalebar= 50µm. **b**) Lungs from Veh or RTX mice were processed for flow cytometry. Cells were gated from CD45 live, singlets. RTX and Vehicle exposed to our pre-exposure and infection model as per Figure 5 had similar SP expression in T-cells (CD3), eosinophils (Siglec F) and mast cells (CD117). N=8 per group. Two-sided t-test. **c**) Mice underwent SP supplementation as per Figure 6a. Cells were stained as per panel b. No differences were demonstrated between groups for SP from leukocytes. N=6-8 per group, One-way ANOVA with Holm-Sidak post-hoc test. Two independent experiments.
